# Perirhinal Cortex Learns A Predictive Map of The Task Environment

**DOI:** 10.1101/2023.03.17.532214

**Authors:** David G. Lee, Caroline A. McLachlan, Ramon Nogueira, Osung Kwon, Alanna E. Carey, Garrett House, Gavin D. Lagani, Danielle LaMay, Stefano Fusi, Jerry L. Chen

## Abstract

Goal-directed tasks involve acquiring an internal model, known as a predictive map, of relevant stimuli and associated outcomes to guide behavior. Here, we identified neural signatures of a predictive map of task behavior in perirhinal cortex (Prh). Mice learned to perform a tactile working memory task by classifying sequential whisker stimuli over multiple training stages. Chemogenetic inactivation demonstrated that Prh is involved in task learning. Chronic two-photon calcium imaging, population analysis, and computational modeling revealed that Prh encodes stimulus features as sensory prediction errors. Prh forms stable stimulus-outcome associations that expand in a retrospective manner and generalize as animals learn new contingencies. Stimulus-outcome associations are linked to prospective network activity encoding possible expected outcomes. This link is mediated by cholinergic signaling to guide task performance, demonstrated by acetylcholine imaging and perturbation. We propose that Prh combines error-driven and map-like properties to acquire a predictive map of learned task behavior.

## INTRODUCTION

The brain generates internal models of the environment that describe the relationship between stimuli, events, and outcomes. Models are learned through experience and can be stored as memories. These memories can be recalled to serve as predictions of upcoming stimuli or outcomes to guide ongoing task behavior. As sensory information is evaluated against internal models, they can generate at least two types of neural signals. Activity can report when sensory information does not match the prediction, referred to as a ‘sensory prediction error’. Activity can also report when sensory information is predictive of an outcome such as reward, referred to as a ‘stimulus-outcome association’. In sensory neocortex, sensory prediction errors are a hallmark of predictive coding, a proposed framework in which predictions of sensory information are generated and evaluated against actual sensory input^1–3^. Stimulus-outcome associations are the basis for cognitive maps in the hippocampus^4–6^, a representation that reduces similar spatial and non-spatial associations to a lower-dimensional ‘abstract’ format^7, 8^. This format is proposed to facilitate generalization of novel stimulus-outcome associations^9, 10^.

The extent to which predictive coding and cognitive maps are aspects of distinct or common neurobiological processes is unclear. Recently, it has been proposed that the two theories could be considered part of a broader framework, referred to as a ‘predictive map’^11, 12^. During goal-directed sensory-guided behavior, sensory prediction errors and stimulus-outcome associations would both be readouts of a single predictive map of the task. This predictive map would be acquired and updated by a combination of error learning to minimize sensory prediction errors and associative learning to strengthen stimulus-outcome associations. The map would be used to predict upcoming task events and infer relationships of novel experiences. Different maps could be flexibly recalled depending on behavioral conditions.

To look for neural evidence of a predictive map, we focused on perirhinal cortex (Prh), a zone of convergence between sensory neocortex and the hippocampus^13–15^. Prh has multiple roles in sensory processing including unitizing features, assigning relational meaning, signaling novelty, and temporal ordering of stimulus items^16–18^. These sensory- and memory-related functions suggest that Prh generates a model of relevant sensory information associated with task behavior. This suggests that functions associated with predictive coding and cognitive maps are combined and expressed in this area. Prh also receives dense cholinergic inputs^19–21^. Acetylcholine is involved in reward expectation and enhancing sensory processing related to predictive coding^22, 23^ as well as memory encoding and retrieval related to cognitive maps^24–26^. Cholinergic signaling could serve as a mechanism that would flexibly establish network states enabling predictive maps to be recalled and utilized in Prh. Here, we investigated whether neural substrates in Prh support the acquisition, representation, and implementation of a predictive map of learned sensory-guided behavior.

## RESULTS

### Perirhinal cortex is necessary for sensory learning tasks

To investigate how predictive maps are acquired and updated, mice were trained to perform a goal-directed task that required them to classify sequentially presented whisker stimuli^27, 28^ (Fig. 1a). A motorized rotor was used to deflect multiple whiskers in either an anterior (A) or posterior (P) direction during an initial ‘sample’ and a later ‘test’ period. Mice were trained to report whether the presented sample and test stimuli were non-matching or matching. In addition to the direction of rotation, deflections were delivered at different speeds (‘fast’ or ‘slow’). Speed can be considered both a second stimulus dimension and a variation in the strength of the rotation direction. This means that animals need to consider relevant (direction) and irrelevant (speed) stimulus features in order to abstract a complex sensory relationship (non-match or match). Temporally dissociating the stimulus features into two trial periods enabled us to investigate how predictive maps are evaluated when features are necessary but not yet sufficient to predict outcome (sample) and when the combined features are abstracted to sufficiently predict the outcome (test).

**Figure 1.**
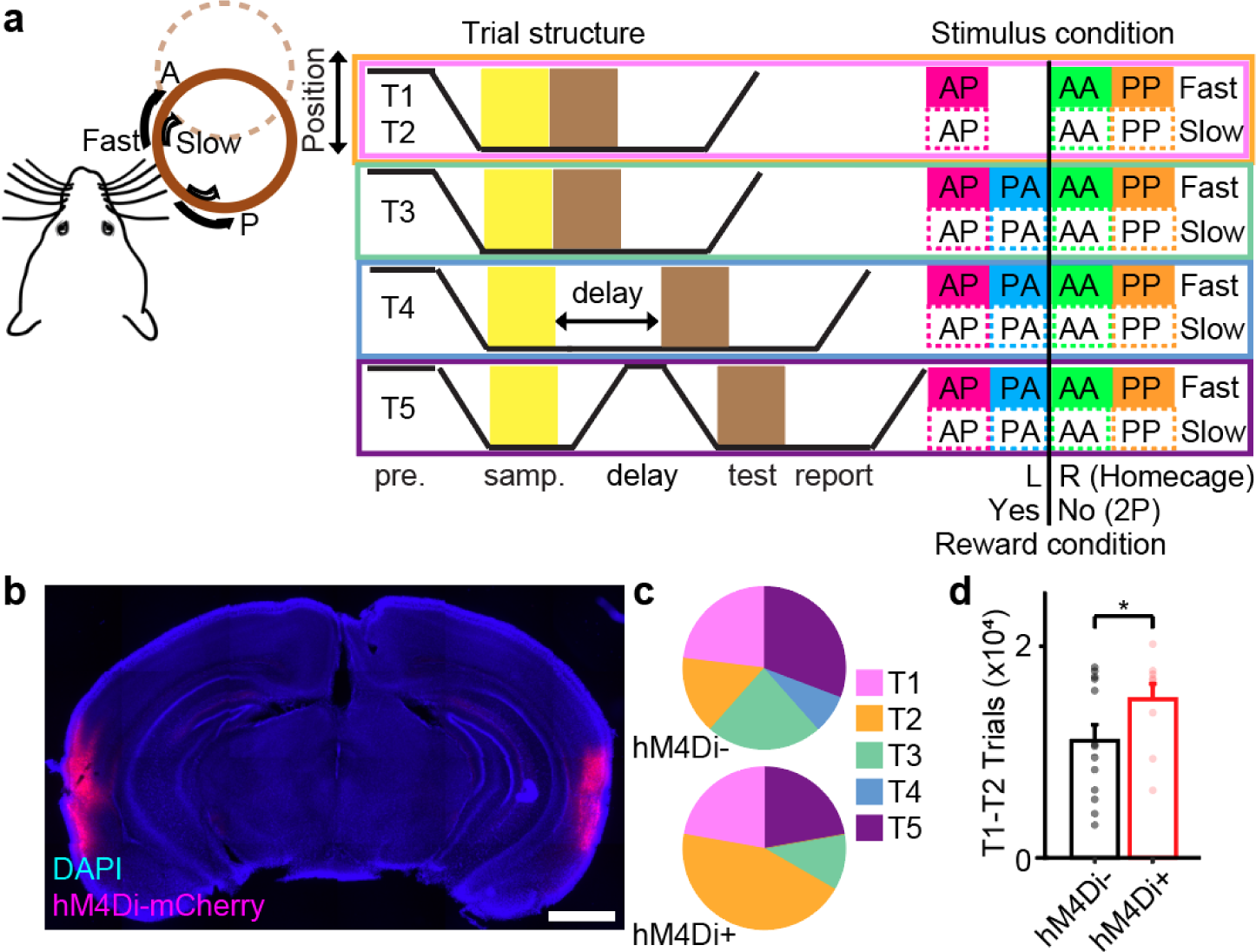
Perirhinal cortex is necessary for learning an abstract sensory task. **a**, Schematic of an abstract sensory learning task. For home cage task training, animal licked left port (L) or right port (R) for reward for non-match or match stimulus conditions, respectively. For head-fixed task training (2P), non-match stimulus conditions were rewarded (Yes) while match conditions were not (No). During head-fixed task training, animals were primarily trained on directions with fast speeds (95% across T1-T4, 75% for T5) with a smaller fraction of slow speeds trials provided as unexpected stimuli (5% across T1-T4, 25% for T5). **b**, Coronal section stained with DAPI (blue) showing bilateral expression of hM4Di-mCherry (magenta) from chemogenetic inactivated animals during home cage task training. **c,** Distribution of final training stage reached for each animal after 84 training sessions for hM4Di-(top) versus hM4Di+ (bottom) groups. The majority of hM4Di+ animals failed to advance past T2. **d,** Number of trials performed in stages T1-T2 by hM4Di-versus hM4Di+ groups. hM4Di+ animals spent more training time in T1-T2. (*\*P*<0.05, Student’s t-test, *n* = 13 hM4Di-animals, 9 hM4Di+ animals). Scale bar = 0.5mm. Error bars = SEM.

Overall training was divided into multiple training stages. Each stage was designed to assay aspects of stimulus-feature and stimulus-reward learning (Table 1-2). The initial training stages consisted of one non-match stimulus condition (AP) and two match stimulus conditions (AA, PP). Training under these conditions was subdivided into 2 stages according to initial naive performance (T1) and learned performance (T2, *d’*>0.45 for two consecutive sessions). Completion of T2 required the animal to unitize the sample and test stimuli and pair it with reward. In the following stage (T3), the remaining held-out non-match condition (PA) was introduced, which required the animal to learn a new stimulus-reward contingency and generalize non-match and match across all possible combinations. Following successful learning of T3, delays between the sample and test stimuli were gradually extended up to 2 seconds (T4) to increase the temporal separation between sample and test stimuli. During the final stage (T5), the rotor was fully retracted during the delay period to require animals to retain a working memory of the sample stimulus. This also prevented the animal from relying on potential positional cues that existed during T4 when the rotor remained in whisker contact throughout the delay period.

**Table 1.** Home-cage task training stages. Summary of task settings utilized at each training stage. Performance criteria indicates the behavioral performance necessary to graduate to the next training stages. NM/M indicates the proportion of stimulus conditions belonging to each category. PA indicates whether that stimulus condition was included in the stimulus set. Fast/Slow indicates the proportion of speed stimulus conditions. Delay indicates the starting and ending delay period length along with the interval in which the delay was increased. Withdraw indicates the distance in which the rotor was withdrawn during the delay period along with the increments of increase.

|  | <u>Performance Criteria</u> | <u>NM/M</u> | <u>PA?</u> | <u>Fast/Slow</u> | <u>Delay (ms)</u> | <u>Withdraw (cm)</u> |
| --- | --- | --- | --- | --- | --- | --- |
| T1 | $d' > 0.45$, 2 sessions | 0.5/0.5 | No | 1/0 | 100ms | 0 |
| T2 | $d' > 1.68$, 2 sessions | 0.5/0.5 | No | 1/0 | 100ms | 0 |
| T3 | $d' > 1.68$, 2 sessions | 0.5/0.5 | Yes | 1/0 | 100ms | 0 |
| T4 | $d' > 1.68$ / 2.05 (skip) | 0.5/0.5 | Yes | 1/0 | 100-2000 (100 inc.) | 0 |
| | $d' > 1.68$ / 2.05 (skip) | 0.5/0.5 | Yes | 1/0 | 200-2000 (200 inc.) | 0 |
| | $d' > 1.68$ / 2.05 (skip) | 0.5/0.5 | Yes | 1/0 | 300-2000 (300 inc.) | 0 |
| | $d' > 1.68$ / 2.05 (skip) | 0.5/0.5 | Yes | 1/0 | 400-2000 (400 inc.) | 0.1-1.5 (0.1 inc.) |
| | $d' > 1.68$ / 2.05 (skip) | 0.5/0.5 | Yes | 1/0 | 500-2000 (500 inc.) | 0.2-1.5 (0.2 inc.) |
| | $d' > 1.68$ / 2.05 (skip) | 0.5/0.5 | Yes | 1/0 | 1000-2000 (500 inc.) | 0.3-1.5 (0.3 inc.) |
| | $d' > 1.68$ / 2.05 (skip) | 0.5/0.5 | Yes | 1/0 | 1500-2000 (500 inc.) | 0.6-1.5 (0.3 inc.) |
| | $d' > 1.68$ / 2.05 (skip) | 0.5/0.5 | Yes | 1/0 | 2000 | 0.9-1.5 (0.3 inc.) |
| | $d' > 1.68$ | 0.5/0.5 | Yes | 1/0 | 2000 | 1.2-1.5 (0.3 inc.) |
| T5 |  | 0.5/0.5 | Yes | 1/0 | 2000 | 1.5 |

**Table 2.** Head-fixed task training stages. Summary of task settings utilized at each training stage. Performance criteria indicates the behavioral performance necessary to graduate to the next training stages. NM/M indicates the proportion of stimulus conditions belonging to each category. PA indicates whether that stimulus condition was included in the stimulus set. Fast/Slow indicates the proportion of speed stimulus conditions. Delay indicates the starting and ending delay period length along with the interval in which the delay was increased. Withdraw indicates the distance in which the rotor was withdrawn during the delay period.

|  | <u>Performance criteria</u> | <u>NM/M</u> | <u>PA</u> | <u>Fast/Slow</u> | <u>Delay (ms)</u> | <u>Withdraw (cm)</u> |
| --- | --- | --- | --- | --- | --- | --- |
| T1 | $d' > 0.45$, 2 sessions | 0.9/0.1 to 0.5/0.5 over 5 sessions | No | 0.95/0.05 | 100 | 0 |
| T2 | $d' > 1.68$, 2 sessions | 0.5/0.5 | No | 0.95/0.05 | 100 | 0 |
| T3 | $d' > 1.68$, 2 sessions | 0.5/0.5 | Yes | 0.95/0.05 | 100 | 0 |
| T4 | $d' > 1.68$ | 0.5/0.5 | Yes | 0.95/0.05 | 100-2000 (100 inc.) | 0 |
| | $d' > 1.68$ | 0.5/0.5 | Yes | 0.95/0.05 | 200-2000 (200 inc.) | 0 |
| | $d' > 1.68$ | 0.5/0.5 | Yes | 0.95/0.05 | 300-2000 (300 inc.) | 0 |
| | $d' > 1.68$ | 0.5/0.5 | Yes | 0.95/0.05 | 400-2000 (400 inc.) | 0 |
| | $d' > 1.68$ | 0.5/0.5 | Yes | 0.95/0.05 | 500-2000 (500 inc.) | 0 |
| | $d' > 1.68$ | 0.5/0.5 | Yes | 0.95/0.05 | 1000-2000 (500 inc.) | 0 |
| | $d' > 1.68$ | 0.5/0.5 | Yes | 0.95/0.05 | 1500-2000 (500 inc.) | 0 |
| | $d' > 1.68$ | 0.5/0.5 | Yes | 0.95/0.05 | 2000 | 1.5 |
| T5 |  | 0.5/0.5 | Yes | 0.75/0.25 | 2000/3000/4000 (0.5/0.25/0.25) prob. | 1.5 |

We first tested whether Prh was required to learn this classification task using chronic chemogenetic inactivation^29^. We utilized a custom-built automated home cage training system that allows for an unbiased assay of task acquisition (**Supplementary Text S1**, Extended Data Fig. 1). Advancement to successive training stages was contingent on pre-defined performance metrics that were applied uniformly to each animal. Reporting of non-match vs. match conditions was carried out by two-alternative forced choice licking of water ports and reinforced by delivery of water reward. We stereotaxically localized whisker-related regions of Prh by first using anatomical tracing to identify sites that exhibit reciprocal connectivity between secondary somatosensory cortex (S2) (Extended Data Fig. 2). Experimental hM4Di+ animals (*n* = 9) received bilateral injections of AAV/9-*hSyn-dio-hM4Di-mCherry* and rAAV-*hSyn-Cre* into the targeted area (Fig. 1b). Control hM4Di-animals (*n* = 13) either received no injection or bilateral sham injections of AAV/9-*hSyn-dio-mCherry* and rAAV-*hSyn-Cre.* All animals were placed in the home cage training system for up to six weeks (∼80 training sessions) with Compound 21 provided in the drinking water to silence neurons in Prh^30^. Histology was performed at the end of behavior experiments to verify viral targeting of Prh. For some animals, the density of hM4Di-mCherry expression (74.9±3.0% of neurons, *n* = 4 animals) along with mRNA *Fos* expression were quantified to verify Prh silencing (Extended Data Fig. 3). Individual hM4Di+ or hM4Di-animals showed a range of learning rates throughout the training period (Extended Data Fig. 4). However, the majority of hM4Di+ animals failed to demonstrate consistent learned behavior to advance past T2 (Fig. 1c). hM4Di+ animals spent more trials in T1-T2 than hM4Di-animals (Fig. 1d, 14,976±1,473 trials hM4Di+ animals vs. 11,058±1,512 trials hM4Di-animals; *P*<0.05, Student’s *t-*test). This demonstrates that Prh is involved in abstract sensory learning.

### Sensory and motor variables across head-fixed task learning

To study how population activity evolves in Prh with task learning, we performed chronic multi-depth two-photon calcium imaging in a separate cohort of head-fixed animals throughout the entirety of training. Virus expressing the genetically encoded calcium indicator, RCaMP1.07 (AAV/PHP.eB*-EF1*α*-RCaMP1.07*), was delivered into Prh. To non-invasively image Prh using an upright two-photon microscope, a 2mm microprism was laterally implanted to provide optical access along the cortical surface using a long working-distance objective (Fig. 2a). Compared to task training in the home cage training systems, task conditions were modified for head-fixed behavior (**Supplementary Text S2**). To help delineate activity between rewarded and non-rewarded stimulus conditions, we employed a go/no-go reward contingency in which only non-match stimulus conditions were rewarded. Compared to the home-cage training, similar performance criteria were applied to advance animals through each stage of training. However, some training parameters were manually tuned to each individual animal to maximize training success (Table 3, **Supplementary Text S3-S4**). Under these conditions, 7 out of 9 animals were successfully trained to T5 within ∼60 training sessions (Fig. 2b). Analysis was performed on animals successfully trained through T5.

**Figure 2.**
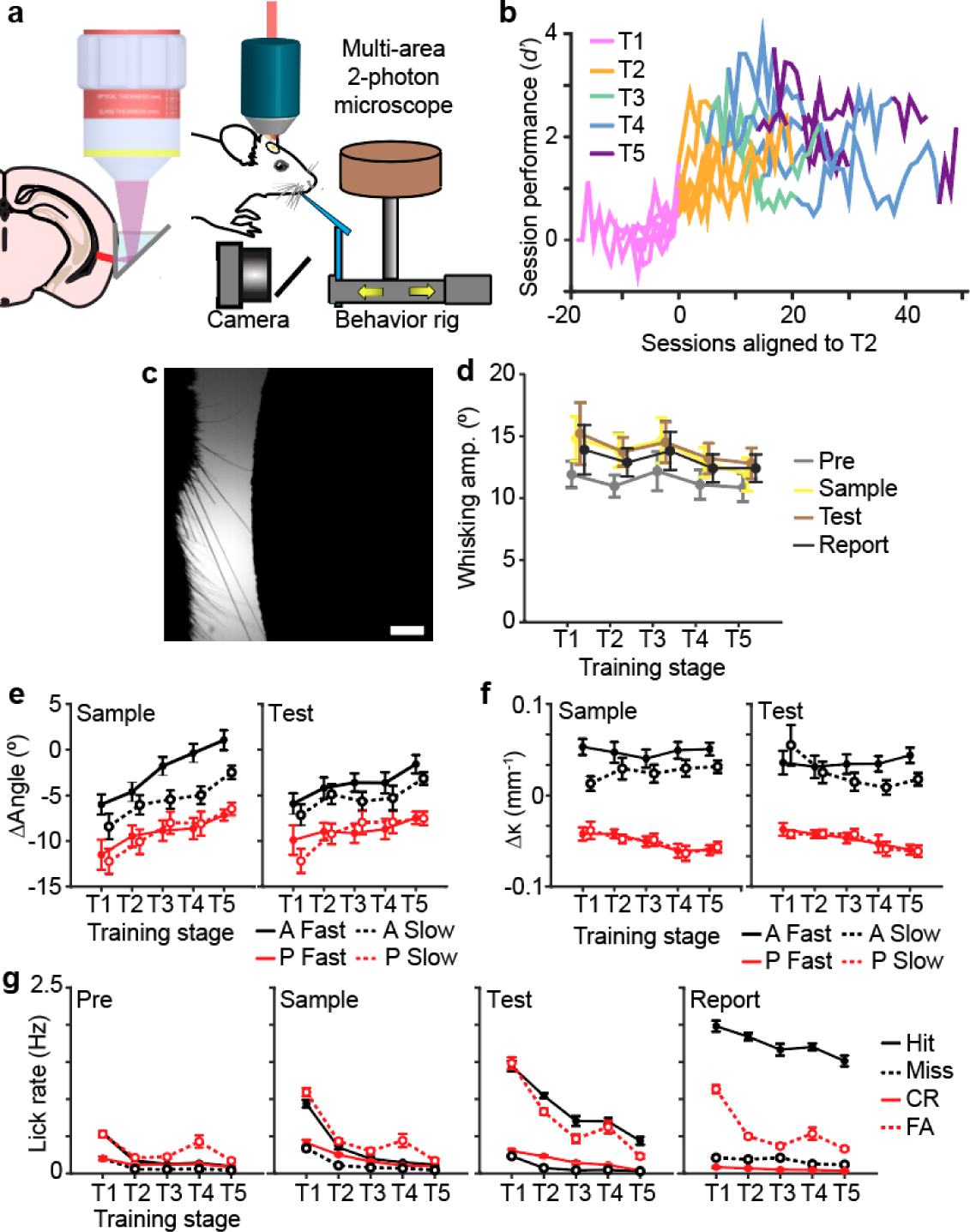
Measuring behavioral correlates throughout task learning. **a,** Schematic of two-photon imaging of Prh using chronically implanted microprisms allowed during head-fixed task training. **b,** Learning curves for individual head-fixed animals trained during two-photon imaging. Only imaged animals reaching T5 were analyzed. **c,** High-speed videography was used to measure whisker kinematics during task behavior. **d,** Whisking amplitude during each trial period across training stages. **e-f,** Change in whisker angle [e] and curvature [f] during sample and test stimulus periods across training stages sorted by speed and direction. **g,** Licking rate during each trial period across training stages sorted by choice. Scale bar = 2mm. Error bars = SEM.

**Table 3.** Training parameters to reinforce correct choice.

| <u>Task</u> | <u>Goal</u> | <u>Criteria</u> | <u>Adjustment</u> |
| --- | --- | --- | --- |
| Head-fixed | Increase punishment to correct for port bias. | *Manual*<br>>70% (hit + false alarm) | *Manual*<br>2-10s time out<br>1-10 air puffs |
| Head-fixed | Decrease punishment to reduce disengagement | *Manual*<br>20-50% miss | *Manual* |
| Home cage | Increase punishment to correct for port bias. | 50 trial sliding window<br><70% correct ($d'$ ~1.05) | Increase 1s time out (10s max)<br>For >7s time out, increase 1 air puff (5 max) |
| Home cage | Decrease punishment to reduce disengagement | 50 trial sliding window<br>>50% miss | Decrease 2s time out and 2 air puffs |
| Head-fixed | Adjust stimulus probability to correct for report biases | *Manual*<br>>70% (hit + false alarm) | *Manual*<br>Up to 0.35/0.65 (NM/M) |
| Home cage | Adjust stimulus probability to correct for report biases | 20 trial sliding window<br>X=% trials favored port<br>Y=% trials neglected port<br>Moderate bias: $X-Y > 0.25$<br>Severe bias: $X-Y > 0.5$ | X=stim. of favored port<br>Y=stim. of neglected port<br>moderate: 0.35/0.65 (X/Y)<br>severe: 0.2/0.8 (X/Y) |
| Both | Adjust stimulus probability to correct for primacy or recency stimulus bias | 20 trial sliding window<br>For non-match stim:<br>X = % correct fav. stim<br>Y = % correct NM stim<br>For match stim:<br>X = % correct fav. stim<br>Y = % correct M stim<br>moderate: $(X/Y - 0.5) > 0.55$<br>severe: $(X/Y - 0.5) > 0.6$ | X=favored stim.<br>Y=neglected stim.<br>moderate: 0.4/0.6 (X/Y)<br>severe: 0.3/0.7 (X/Y) |

We first asked whether animals changed their behavioral strategies with learning by measuring changes in sensory or motor variables. In addition to two-photon calcium imaging, high-speed videography was performed to measure whisker kinematics and whisking behavior (Fig. 2c, Extended Data Fig. 5). Unlike in other whisker-based sensory tasks^31, 32^, animals did not actively whisk during task performance. Whisking amplitude did not significantly change across training stages (Fig. 2d). We additionally examined licking behavior across training. In early training stages, animals showed sporadic licking across different trial epochs such as the sample and test period, but this became more restricted to the report period as animals advanced in the task (Fig. 2g, pre, *P*<1×10^−29^, *F_4,1159_*= 38.8; sample, *P*<1×10^−78^, *F_4,1159_*= 109.0; test, *P*<1×10^−43^, *F_4,1159_*= 57.2; report, *P*<1×10^−5^, *F_4,1159_*= 8.3, one-way ANOVA with post-hoc multiple comparison test).

We next compared whisker kinematics during different direction and speed conditions. Overall, whisker angle changes trended more in the anterior direction (Fig. 2e**, sample**, *P*<1×10^− 5^, *F_4,593_* = 8.01, one-way ANOVA with post-hoc multiple comparison test; **test**, *P*<1×10^−4^, *F_4,592_* = 7.10, one-way ANOVA with post-hoc multiple comparison test). Despite this, posterior stimuli consistently produced more negative angle deflections than anterior stimuli. Posterior stimuli also consistently produced more negative curvature changes than anterior stimuli (Fig. 2f). Compared to fast conditions, slow conditions produced weaker negative angle deflections and curvature changes in the anterior direction. No difference was observed for either angle or curvature changes between slow and fast stimuli in the posterior direction.

### Perirhinal cortex learns sensory prediction errors

Given the specific changes in sensory and motor variables across learning, we sought to determine what aspects of sensory information are encoded in Prh. We focused on neural activity related to stimulus direction or speed and its relationship to task performance. Animals were primarily trained on directions with fast speeds (95% across T1-T4, 75% for T5) with slow speed trials provided as less frequent stimuli (5% across T1-T4, 25% for T5). Since whisker kinematic analysis shows that slower speeds produce less deflections in the anterior direction, weaker information about stimulus direction could affect task performance on slow speed trials. Indeed, while animals were able to learn the task at fast and slow speeds, they performed worse on slow compared to fast speed conditions as they approached later training stages (T4, *P*<0.05; T5, *P*<0.05, paired Student’s *t*-test, Fig. 3a).

**Figure 3.**
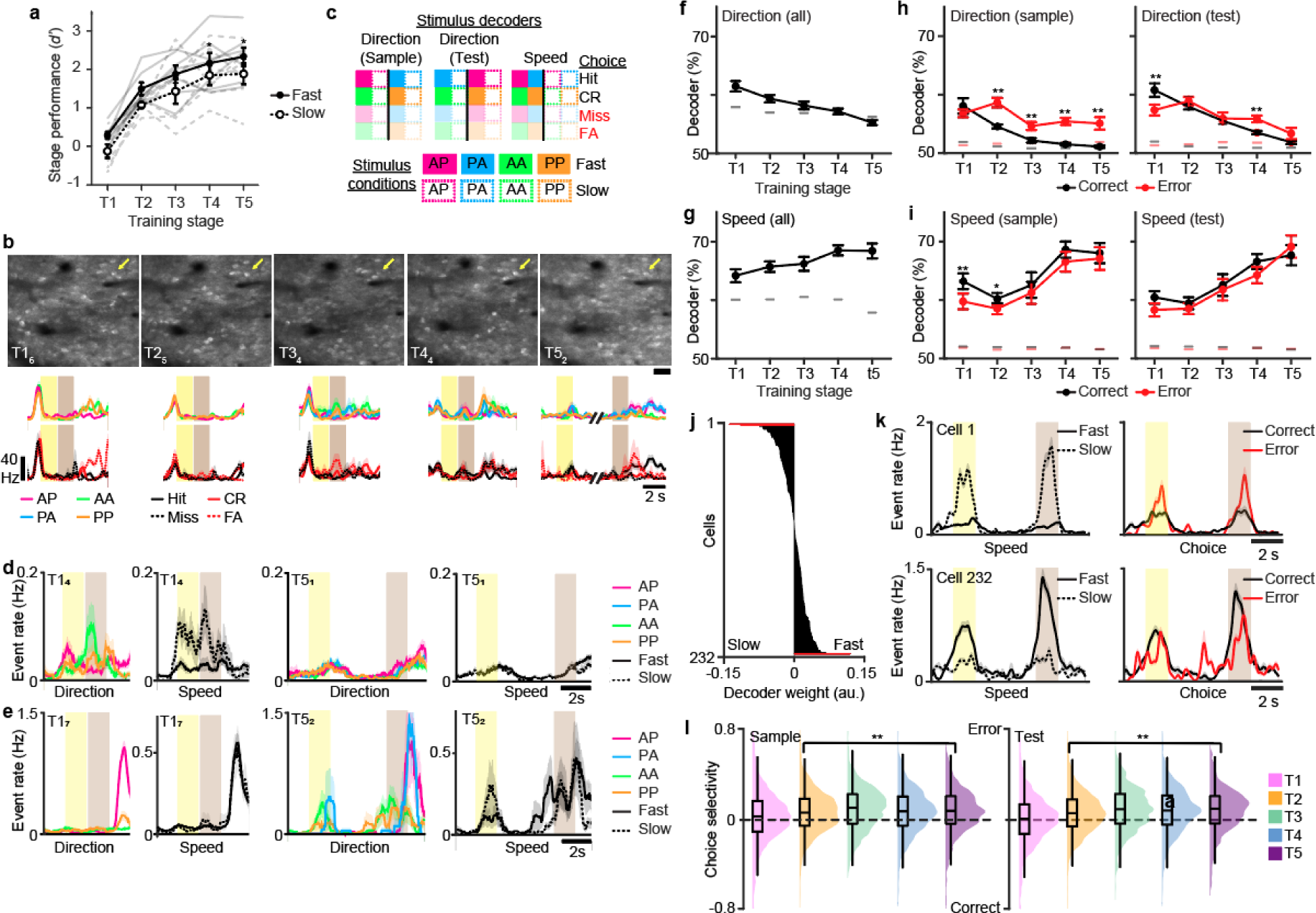
Perirhinal cortex learns sensory prediction errors. **a,** Behavioral performance across training stages separated by fast versus slow speed trials. **b,** Example imaging area at denoted training stage and session number (top row). Mean activity sorted by stimulus condition or choice (bottom row) for indicated neuron (yellow arrow). **c,** Schematic of population decoders to stimulus direction or speed. Black line separates decoder trial types. For correct trials, only hit and correct rejection (CR) trials were used. For error trials, only miss and false alarm (FA) trials were used. **d,** Example neuron with selectivity to direction and speed during early training sessions (T1_4_) that showing reduced selectivity in expert sessions (T5_1_). **e,** Example neuron with developing selectivity to speed in expert sessions (T5_2_). **f,** Decoder performance to stimulus direction across training stages (*P* < 1×10^−8^, one-way ANOVA with post-hoc multiple comparison test). **g,** Decoder performance to stimulus speed across training stages (*P*<0.02, one-way ANOVA with post-hoc multiple comparison test). **h-i,** Decoder performance to stimulus direction [h] or speed [i] across training stages during the sample (left) and test (right) stimulus period separated by correct versus error trials (Student’s *t*-test). **j,** Example population vector weights for decoder to stimulus speed from one imaging session. Significant weights are indicated (red). **k,** Mean event rates for example neurons with significant weights in [j] sorted by fast versus slow speed trials (left) or correct versus error trials (right). **l,** Distribution and box plot of choice selectivity during sample (left) or test (right) stimulus period for speed-tuned neurons across training stages (sample period: *P*<1×10^−15^; test period: *P*<1×10^−41^, one-way ANOVA with post hoc multiple comparison test). Lines indicate 95^th^ percentile of shuffled performance in [f-i]. Error bars = SEM; [f-i]. \*\**P*<0.005 for [f-i]. *n* = 70 T1 sessions, 75 T2 sessions, 30 T3 sessions, 79 T4 sessions, 48 T5 sessions from 7 animals for [f-i]. *n* = 529 neurons from 7 animals for [l].

We analyzed how Prh encodes direction and speed across training. For every training session, neuronal populations (*n* = 2335 neurons, 7 animals) in layer 2/3 (L2/3) of Prh were simultaneously imaged across 2 imaging depths using a multi-area two-photon microscope (Fig. 3b, Extended Data Fig. 6)^33^. In single cells, we observed examples of preferred responses to stimulus direction during early training sessions that disappeared in later sessions (Fig. 3d). We also observed selectivity to stimulus speed emerging over training sessions (Fig. 3e). To characterize these changes at a population level, population decoding was performed on trial conditions related to direction or speed (Fig. 3c). Early during learning, direction could be decoded above chance but gradually decreased to chance levels by T5 (Fig. 3f, *P*<1×10^−8^, *F_4,266_* = 12.65, one-way ANOVA with post-hoc multiple comparison test). In contrast, decoders trained to speed increased performance with learning (Fig. 3g, *P*<0.02, *F_4,262_* = 3.19, one-way ANOVA with post-hoc multiple comparison test). Overall, this indicates that task training results in weakening representations of task relevant stimuli (direction) and strengthening of task irrelevant stimuli (speed) in Prh.

The above results are in opposition to previous results observed in primary somatosensory cortex (S1) during task learning which is typified by strengthening of task relevant features^32, 34^. They are also inconsistent with the changes in whisker kinematics observed across training stages in the high-speed videography. We assessed whether activity related to direction or speed differed depending on the animals’ choice. Decoders were trained on direction or speed separately for correct (‘hit’ or ‘correct rejection’) or error (‘miss’ or ‘false alarm’) trials. For direction, we found that decoder accuracy during the sample period decreased to chance over learning on correct trials, but this information remained above chance on error trials (Fig. 3h). In contrast, analysis of previously acquired S1 population data in expert animals performing the task showed that direction was stronger on correct compared to error trials (Extended Data Fig. 7). In Prh, decoder performance for speed increased similarly for correct and error trials (Fig. 3i). To more closely examine how speed selectivity relates to choice selectivity in single neurons, we identified neurons with significant population decoder weights to speed (Fig. 3j). We then compared the firing rates of these neurons when sorted for speed conditions versus correct choice conditions. We found examples of neurons that were tuned to both speed and choice (Fig. 3k). We measured the choice-selective response distribution of speed-tuned neurons across learning. While the distribution of speed-tuned neurons showed balanced responses to choice during T1, choice selectivity became biased towards error trials once animals demonstrated learned performance (T2-T5) (Fig. 3l, sample: *P*<1×10^−15^, *F_4,7578_*=19.69, one-way ANOVA with post-hoc multiple comparison test; test: *P*<1×10^−41^, *F_4,7682_* =50.69, one-way ANOVA with post-hoc multiple comparison test).

These neural signatures can possibly be explained by Prh’s role in familiarity and novel object recognition^35^. Familiarity can be detected by comparing, through subtraction, the current sensory input to one that was previously stored in memory^36^. As sensory information is stored into memory, subtraction results in reduced responses for familiar stimuli and increased responses for novel stimuli. A similar mechanism could be employed for encoding direction and speed during task learning. Memories of direction, as a task-relevant stimuli, may be preferentially stored instead of speed in connected brain areas such that only that component will be subtracted from the current stimulus when compared in Prh. To illustrate this, we constructed a simple model, focusing on encoding the stimulus features while neglecting models involving working memory^37^ or comparison of match and non-match^38^ which have been explained previously. The model consists of an autoencoder with input, hidden, and output layers analogous to S1, the hippocampus, and Prh, respectively. The input to the model consists of two stimulus dimensions corresponding to direction and speed (Fig. 4a, left). The network was trained to reconstruct the input in the output layer. An additional output neuron was trained to generate the correct response required to get reward. This neuron biased the representation of the hidden layer of the autoencoder to make the direction of motion more relevant than speed. We also limited the activity in the hidden layer by imposing a sparseness constraint (L2-norm) (Fig. 4a, right; **see Methods**). Finally, a downstream neuron read out the familiarity signal, that is, the difference between the reconstructed output and input^36^. With these simple components, we were able to reproduce the experimental results. Information about the task-relevant variable direction of motion decreased, whereas information about speed increased throughout learning (Fig. 4b). Importantly, this result was only possible when all components were included in the model (see Extended Data Fig. 8).

**Figure 4.**
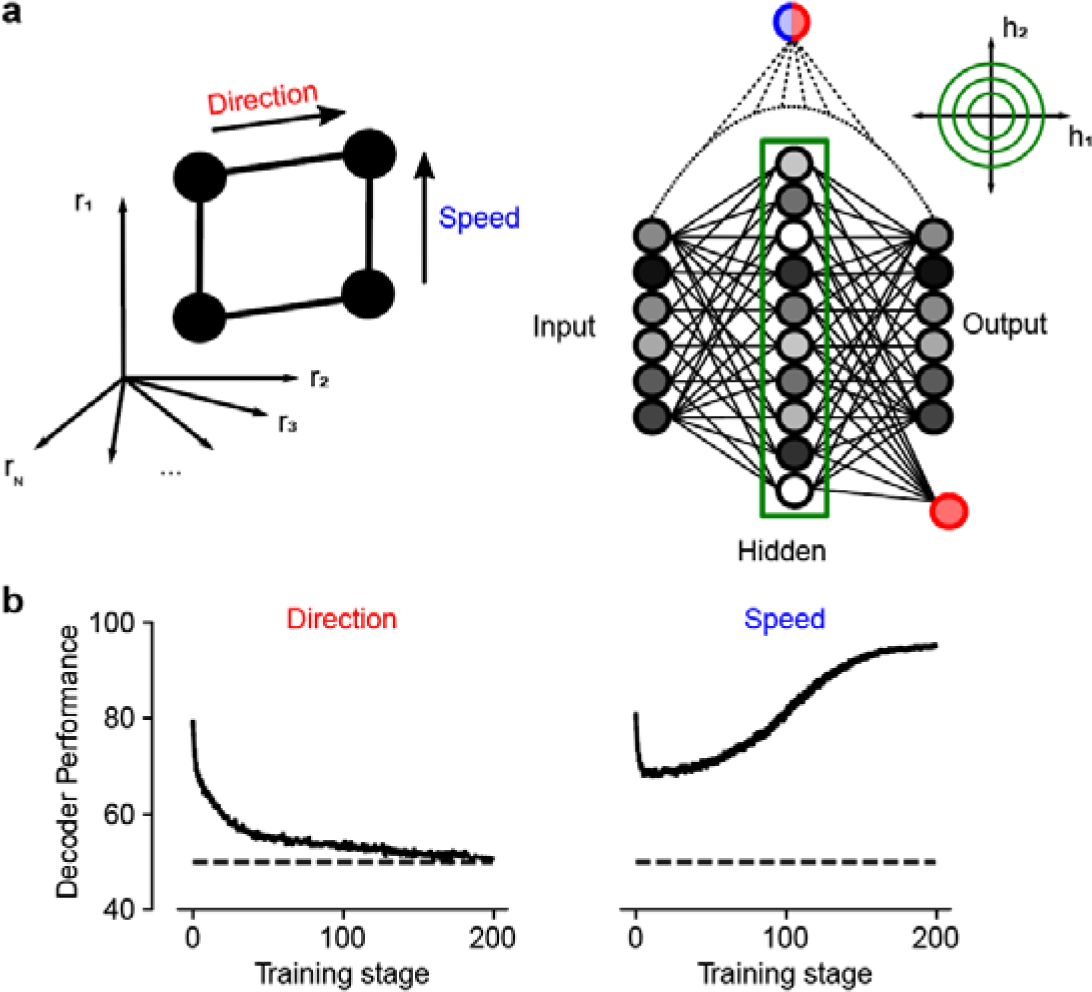
Computational model of sensory prediction errors in perirhinal cortex. **a,** An autoencoder with three layers (input, hidden, and output) was trained to represent the input. The input consisted of two independent stimulus variables: direction of motion (red) and speed (blue). A downstream neuron was trained (logistic regression) to decode direction (red) and speed (blue) by reading out the difference between the reconstructed output and the input (dotted line). Sparsity in the hidden layer was imposed by adding an L2-norm term on the loss function. **b,** Decoding performance of direction (red, left) and speed (right, blue) as a function of training epoch for the downstream neuron reading out from familiarity activity. Similar to experimental results, decoding performance of direction decreases, whereas decoding performance for speed increases throughout training. Error bars correspond to SEM across independent simulations (*n* = 50). See also Extended Data Fig. 8.

Overall, the results above indicate the Prh does not represent sensory information in the same manner as S1 does. Instead, it suggests that stimulus activity in Prh may reflect a sensory prediction error signal (ie. the difference between actual and expected sensory information), consistent with theories of predictive coding^3^ and Prh’s role in familiarity and novel object recognition. Information about direction decreases as Prh forms an internal model of direction as the task relevant feature, explaining away the delivered stimuli. Concurrently, information about speed increases to signal the prediction error between directions that are presented at the expected fast speeds versus the unexpected, weak slow speeds.

### Stimulus-reward associations emerge and stabilize with learning

To understand how sensory and reward information are integrated to form stimulus-reward associations, we analyzed how representations of reward outcome evolved across learning. A cross-session decoder was trained using Hit vs. non-Hit trials from one session and tested on other sessions across learning (Fig. 5a). When assessing cross-session performance between neighboring sessions during the report period, representations of reward outcome were stably represented above chance on a session-to-session basis. No differences in session-to-session performance were found between training stages (Fig. 5b, *P*=0.19, *F_4,260_*= 1.54, one-way ANOVA). Analysis of cross-session performance across longer time scales and across training stages showed that representations of reward outcome were less stable early in training (T1) but stabilized as animals learned the task (Fig. 5c). These results suggest that learning produces a stable, long-term representation of reward outcome.

**Figure 5.**
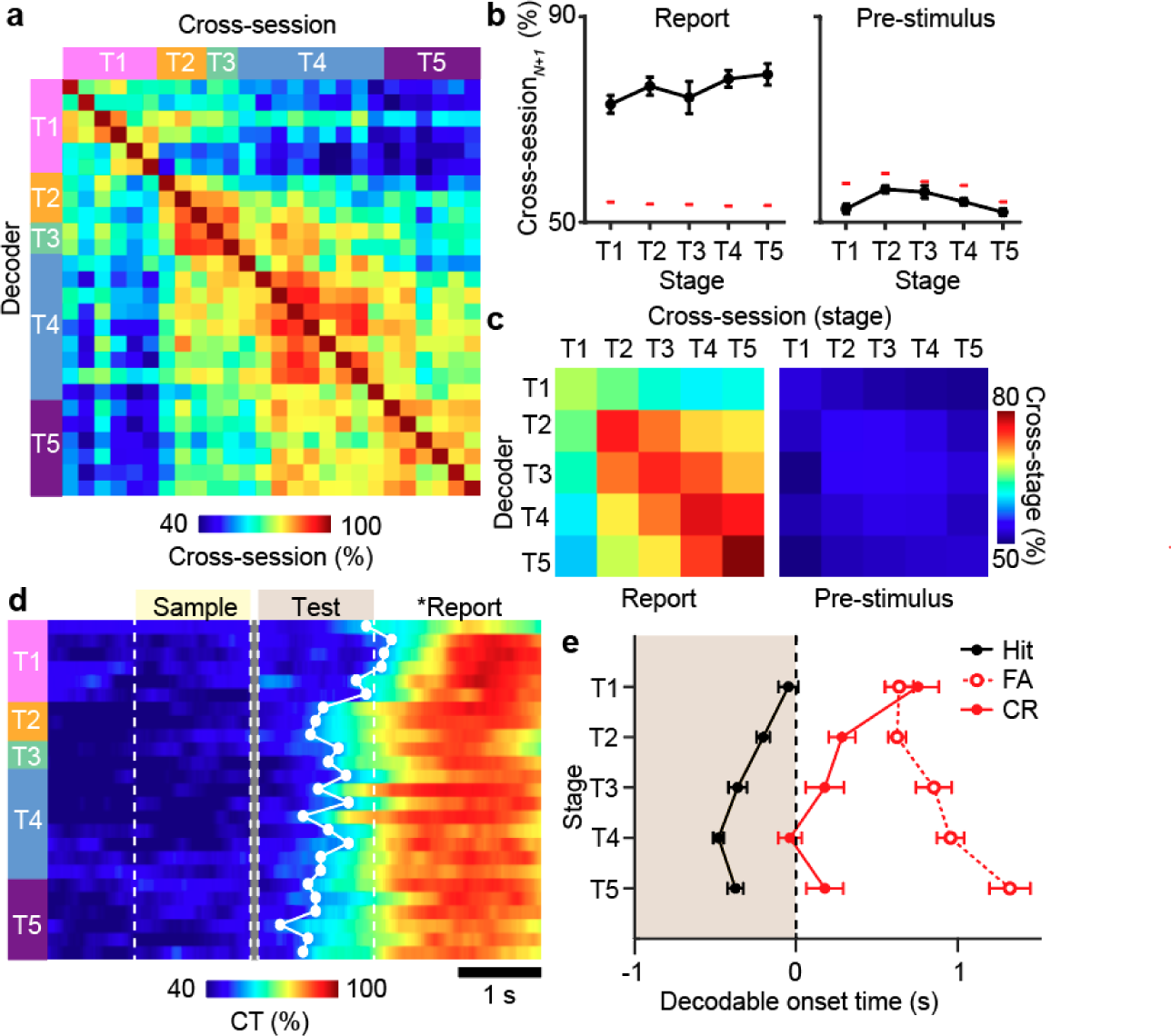
Learning of stimulus-reward associations. **a,** Example of cross-session decoder performance trained on Hit trials during the report period for one animal across training. **b,** Cross-session performance for decoders trained on session*_N_* and tested on session*_N_*_+1_ for report activity (left) or pre-stimulus activity (right) across training stages. **c,** Cross-session decoder performance across training stages for report activity (left) or pre-stimulus activity (right). **d,** Example of cross-temporal (CT) decoder for reward conditions trained on report activity across each training session for one animal. First decodable time point above chance is shown (white dot). **e,** Decodable onset timepoint for cross-temporal decoder of report activity for decoders trained on hit, false alarm, or correct rejection trials (*P*<0.002, *F_4,282_*=4.44, one-way ANOVA with post-hoc multiple comparison test. Error bars = SEM. Lines indicate 95^th^ percentile of shuffled performance [b]. *n* = 70 T1 sessions, 75 T2 sessions, 30 T3 sessions, 79 T4 sessions, 48 T5 sessions from 7 animals.

Given that reward outcome stabilizes with learning, we asked whether such representations reflect a stimulus-reward association which would precede reward delivery. A cross-temporal decoder was trained on Hit. vs non-Hit trials during the report period and then tested on time points across the trial period. We identified a gradual retrograde expansion of decoder performance related to reward outcome over the course of learning that preceded reward and extended into the test stimulus period (Fig. 5d). Analysis of the onset of decodable reward outcome across training stages showed that this expansion emerged as animals demonstrated learned performance (T2) and continued to expand throughout the additional training stages (Fig. 5e, *P*<0.002, *F_4,282_*=4.44, one-way ANOVA with post-hoc multiple comparison test). To test whether this temporal expansion is specific to rewarded trials, we conducted similar analysis of cross-temporal decoders trained to non-rewarded conditions that controlled for either licking behavior (false alarm) or correct choice (correct rejection). Neither decoder showed onset accuracy that extended into the test period. This demonstrates that neural representations on Hit trials correspond to a stimulus-reward association. The temporal profile of this expansion suggests that this association emerges in a retrograde manner from reward outcome.

### Stimulus-reward associations generalize in an abstract format

We next asked whether stimulus-reward associations were specific to a given stimulus set or could generalize across stimulus conditions. To address this, we analyzed how representations changed from T2 to T3 when the novel PA stimulus-reward contingency was introduced. Behaviorally, mice were flexibly able to correctly respond on the first session in which PA was introduced (T3_0_). Performance on PA further improved over ∼4-5 sessions, reaching similar levels as AP (Fig. 6a). We observed examples of single cells that exhibited distinct temporal responses between AP and PA conditions at T3_0_. These responses changed over sessions, resulting in similar responses between the two conditions (Fig. 6b). To characterize these changes at a population level, we trained two separate population decoders on activity during the report period on rewarded conditions using either only AP or PA (Fig. 6c). This allowed us to independently evaluate each representation across T3 sessions. Cross-temporal analysis showed that the temporal profile of AP and PA representations were distinct at T3_0_ but became similar after 4 sessions (T3_4_) (Fig. 6d). Whereas the onset accuracy extended into the test period for AP at T3_0_, indicative of a stimulus-reward association, onset accuracy for PA initially was restricted to the report period but expanded into the test period over the course of 3-4 sessions (Fig. 6e, *P*<0.002, *F_9,54_*=3.64, two-way repeated measures ANOVA with post-hoc Student’s *t-*test). This demonstrates that acquisition of new stimulus-reward contingencies occurs through a common mechanism of retrograde expansion from reward outcome.

**Figure 6.**
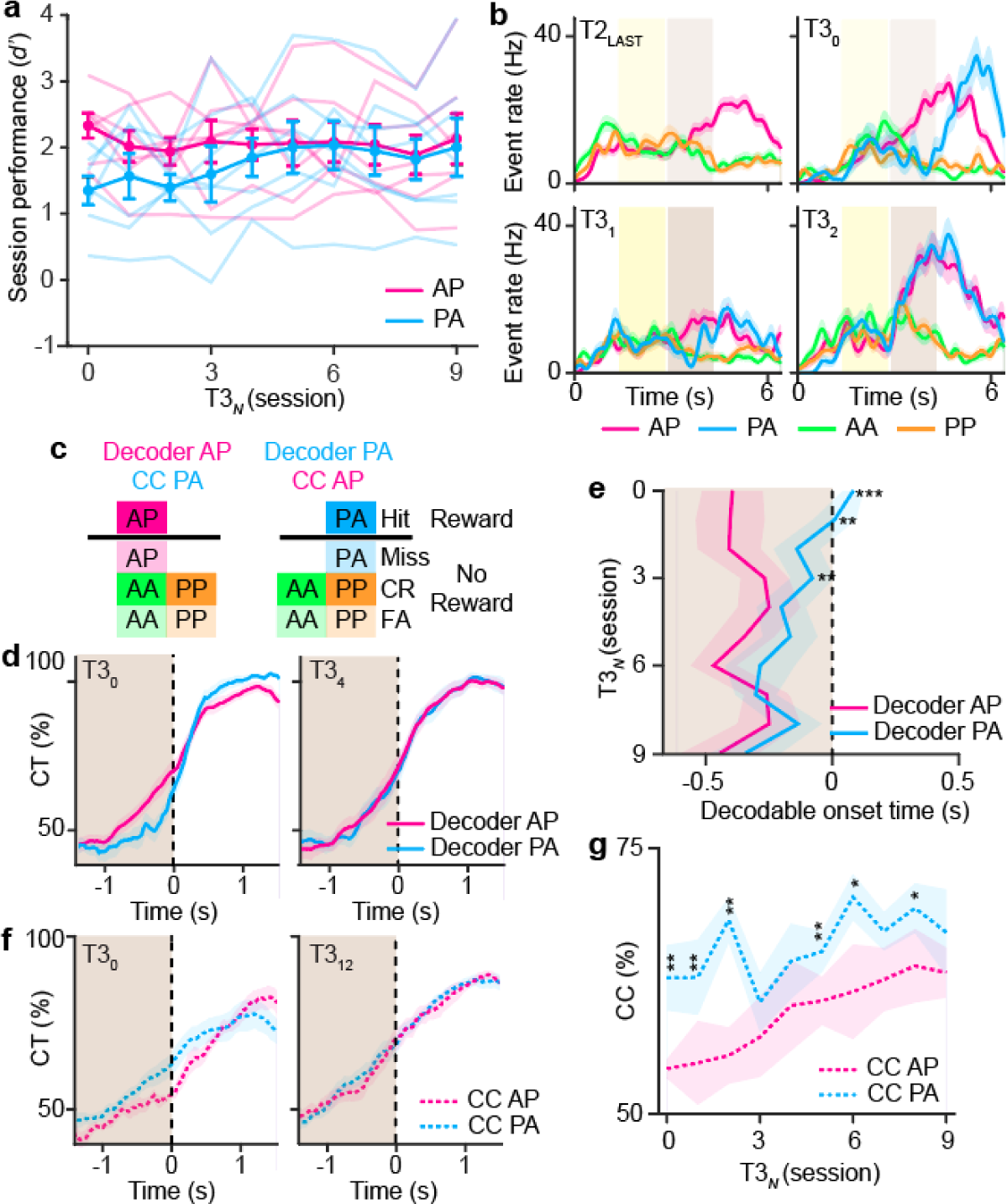
Stimulus-reward associations are abstract. **a,** Behavioral performance aligned to the first T3 session for AP versus PA stimulus conditions. Mean and individual animal performance is shown. **b,** Mean activity in an example neuron separated by stimulus conditions across the first four T3 sessions. **c,** Schematic for population decoder for reward using either only AP or PA stimulus conditions. Cross-condition (CC) decoder also shown for the complementary condition. **d,** Cross-temporal decoder performance trained on report activity for the rewarded AP or PA condition during the T3_0_ or T3_4_ session. **e,** Decodable onset timepoint for either the rewarded AP or PA condition T3 sessions (*P*<0.002, two-way repeated measures ANOVA with post-hoc Student’s *t-*test). **f,** Cross-temporal decoder performance trained on report activity for the rewarded AP or PA condition and tested on the cross condition during the T3_0_ or T3_12_ session. **g,** Cross-temporal decoder performance trained on report activity for the rewarded AP or PA condition and tested on the cross condition test period activity across T3 sessions (*P*<0.05). Error bars = SEM. \**P*<0.05, \*\**P*<0.02, \*\*\**P*<0.001 for [e] and [g]. *n* = 7 animals for [b, d-g].

Representations of AP-reward and PA-reward associations could exist in different or similar neural subspaces. The latter would imply that the geometry of stimulus-reward associations in Prh are represented in an abstract format^7^. To test this, we analyzed the cross-condition performance for each of the two separate population decoders (ie. testing AP performance using a PA decoder and vice-versa). Cross-condition PA performance to the AP decoder during the test stimulus period was initially worse than the opposite cross-condition but gradually improved over the course of 9 sessions (Fig. 6f,g, *P*<0.05, *F_9,54_*=2.16, two-way repeated measures ANOVA with post-hoc Student’s *t-*test). This suggests acquisition of new stimulus-reward contingences occurs in two phases: an initial establishment of the stimulus-reward association followed by a consolidation that aligns the new association into the same subspace of existing stimulus-reward associations. Overall, these findings demonstrate that Prh can generalize across novel stimulus-reward contingencies to form stimulus-reward associations that are representationally abstract.

### Neural signatures of expected outcome in Prh

The observation that stimulus-outcome associations emerge in a retrograde manner to precede the report period suggests that stimulus information is integrated with ongoing activity that could signal an expected outcome (ie. reward delivery). Neural activity reflecting the expectation of reward or punishment has been observed during task engagement in other brain areas^39^. Therefore, we asked whether ongoing Prh activity throughout the trial period could contain an expectation signal of future outcomes. We looked for evidence of population activity corresponding to expected outcome. This was defined by the ability for a linear decoder to decode trial outcome when trained on activity at the beginning of the trial during the pre-stimulus period. Additionally, we asked whether this population subspace was maintained across the trial epoch to link task events to a given outcome. This was defined as the ability for the same decoder to cross-temporally decode trial outcome when tested on activity during the report period.

Two separate population decoders were trained on either hit vs. non-hit trials (Expected Hit) or correct rejection (CR) vs. non-CR trials (Expected CR) during the pre-stimulus period (Fig. 7a). When trained and tested during the pre-stimulus period (Fig. 7b), trial outcome could be decoded above chance throughout training. The accuracy of this decoder was consistently weaker than decoders trained and tested during the report period (Fig. 7d). Cross-session decoders to Expected Hit were not able to perform above chance, suggesting that this activity is unstable across sessions unlike the stimulus-reward associations (Fig. 5c). Expected Hit could also be decoded during the sample and test period (Extended Data Fig. 9). These same decoders did not encode information about stimulus direction or speed indicating that expected outcome activity occupied a different subspace from sensory prediction errors.

**Figure 7.**
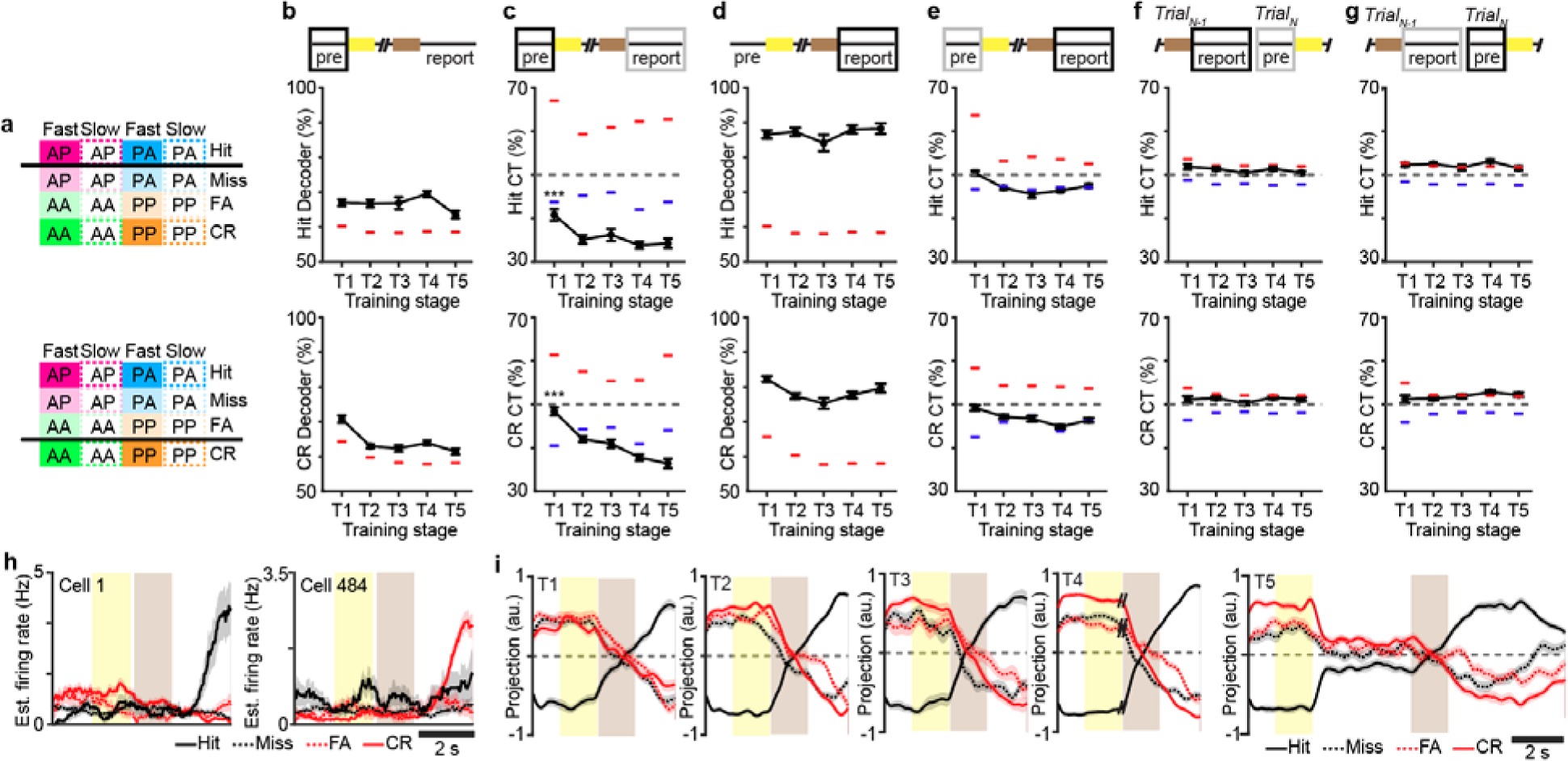
Perirhinal cortex encodes expected outcome throughout task learning. **a,** Schematic of population decoder trained to Expected Hit (top) or Expected CR (bottom). **b,** Decoder performance to Expected Hit (top) and Expected CR (bottom). **c,** Cross-temporal (CT) decoder performance to Expected Hit (top) and Expected CR (bottom). Black box indicates trained time window during the pre-stimulus period. Grey box indicates (solid box) tested time window during the report period. **d,** Decoder performance during the report period for Hit (top) and CR (bottom). **e,** CT decoder performance trained during the report period (black box) and tested during the pre-stimulus period (grey box) for Hit (top) and CR (bottom) trials. **f,** CT decoder performance trained during the report period (black box) and tested during the pre-stimulus period of the following trial (grey box) for Hit (top) and CR (bottom) trials. **g,** CT decoder performance trained during the pre-stimulus period (black box) and tested during the report period of the previous trial (grey box) for Hit (top) and CR (bottom) trials. **h,** Mean estimated firing rate for example neurons with significant weights for Expected Hit decoder. Cell 1 shows elevated firing during the pre-stimulus period on CR trials but strongly responds during the report period of Hit trials. Cell 484 shows elevated firing during the pre-stimulus period on Hit trials but strongly responds during the report period of CR trials. **i,** Projection of neural activity along the decision variable for Expected Hit [c] across the trial period sorted by trial type across training stages. Error bars = SEM., \*\*\**P*<1×10^−3^, one-way ANOVA with post-hoc multiple comparisons test. For [b-g], dashed lines indicate 95^th^ percentile (red) and 5^th^ percentile (blue) of nulled performance for classifier after shuffling trial labels. *n* = 70 T1 sessions, 75 T2 sessions, 30 T3 sessions, 79 T4 sessions, 48 T5 sessions from 7 animals.

Analysis also demonstrates that this subspace is maintained throughout the trial period. Decoders trained on the pre-stimulus period were able to decode outcome activity below chance when tested during the report period (ie. below the 5^th^ percentile of the shuffled distribution) (Fig. 7c). This was particularly strong during T2-T5 sessions when animals exhibited strong task performance (Expected Hit: *P*<1×10^−5^, *F_4,296_*=7.64; Expected CR: *P*<1×10^−19^, *F_4,285_*=29.18, one-way ANOVA with post-hoc multiple comparisons test). To better understand how pre-stimulus activity predicts outcome activity below chance in single neurons, we identified neurons with significant population decoder weights. These neurons exhibit low levels of activity during the pre-stimulus period that differed slightly when sorted between Hit, Miss, FA, and CR trials. One neuron that showed slightly elevated pre-stimulus activity on CR trials showed robust outcome responses on Hit trials. Another neuron that showed slightly elevated pre-stimulus activity on Hit trials showed robust outcome responses on CR trials (Fig. 7h). We examined the population trajectory along the subspace of the pre-stimulus decoder (Fig. 7i). For Expected Hit, the population activity was projected along the decision variable axis for each of the 4 choice conditions over the time course of the trial. We observed that activity on hit and non-hit trials was separated along the axis through the pre-stimulus and sample stimulus period. The trajectories converged during the test stimulus period and then flipped their sign during the report period. This suggests that the decoder trained on expected outcomes captures neurons whose firing intially favors one potential trial outcome during the pre-stimulus period but later reverses its response to prefer the actual outcome during report period. The sign flip along this subspace explains the below chance performance during the report period.

To confirm that activity in the pre-stimulus period constitute a prospective and not a retrospective signal, we analyzed the performance of several cross-temporal decoders. Cross-temporal decoder trained during the report period was not able to explain reward information during the pre-stimulus period (Fig. 7e). To test if pre-stimulus information reflects a trial history of recent outcomes as observed in other cortical areas^40^, cross-temporal decoders between the pre-stimulus and the report period of the previous trial were tested (Fig. 7f,g). These decoders did not perform above chance. Overall, this demonstrates that activity early in the trial constitutes a prospective signal whose subspaces emerges with training to link expectation to learned outcomes.

### Cholinergic signaling mediates expected outcome calcium signals

To investigate how expected outcome signals are established in Prh and whether they link expectations with outcome in a behavior-dependent manner, we looked for signaling mechanisms that could mediate these neural dynamics. Acetylcholine (Ach) is a major neuromodulator that affects the state of cortical networks^22^ and has been associated with reward expectation^41^. We hypothesized that Ach signaling could establish expected outcome states in Prh. To visualize Ach activity during early stages of training (T1 and T2), we imaged Ach release in Prh using the fluorescent Ach indicator GRAB-Ach3.0^42^ (Fig. 8a). Bulk Ach signals were measured across the field of view. Prominent high Ach release was measured during the pre-stimulus period across all trials (Fig. 8b,c, Extended Data Fig. 10). On hit trials, increases in Ach were also observed to be related to licking behavior prior to reward delivery but not during reward consumption. Similar dynamics were observed on false alarm trials when no reward was delivered. These dynamics suggest that Ach in Prh signals behavioral correlates of reward expectation. To quantify the relationship between Ach and the behavioral task, we modeled Ach signals using a generalized linear model (GLM) with task variables representing the pre-stimulus period, stimulus direction, pre-reward licking, post-reward licking, reward delivery, and the post-trial period (Fig. 8d,e, Extended Data Fig. 11). The pre-stimulus task variable best explained Ach signals and increased from T1 to T2 (Fig. 8f, *P*<0.05, Student’s *t*-test). This increase in pre-stimulus Ach coincided with the emergence of sustained expected outcome signals (Fig. 7c).

**Figure 8.**
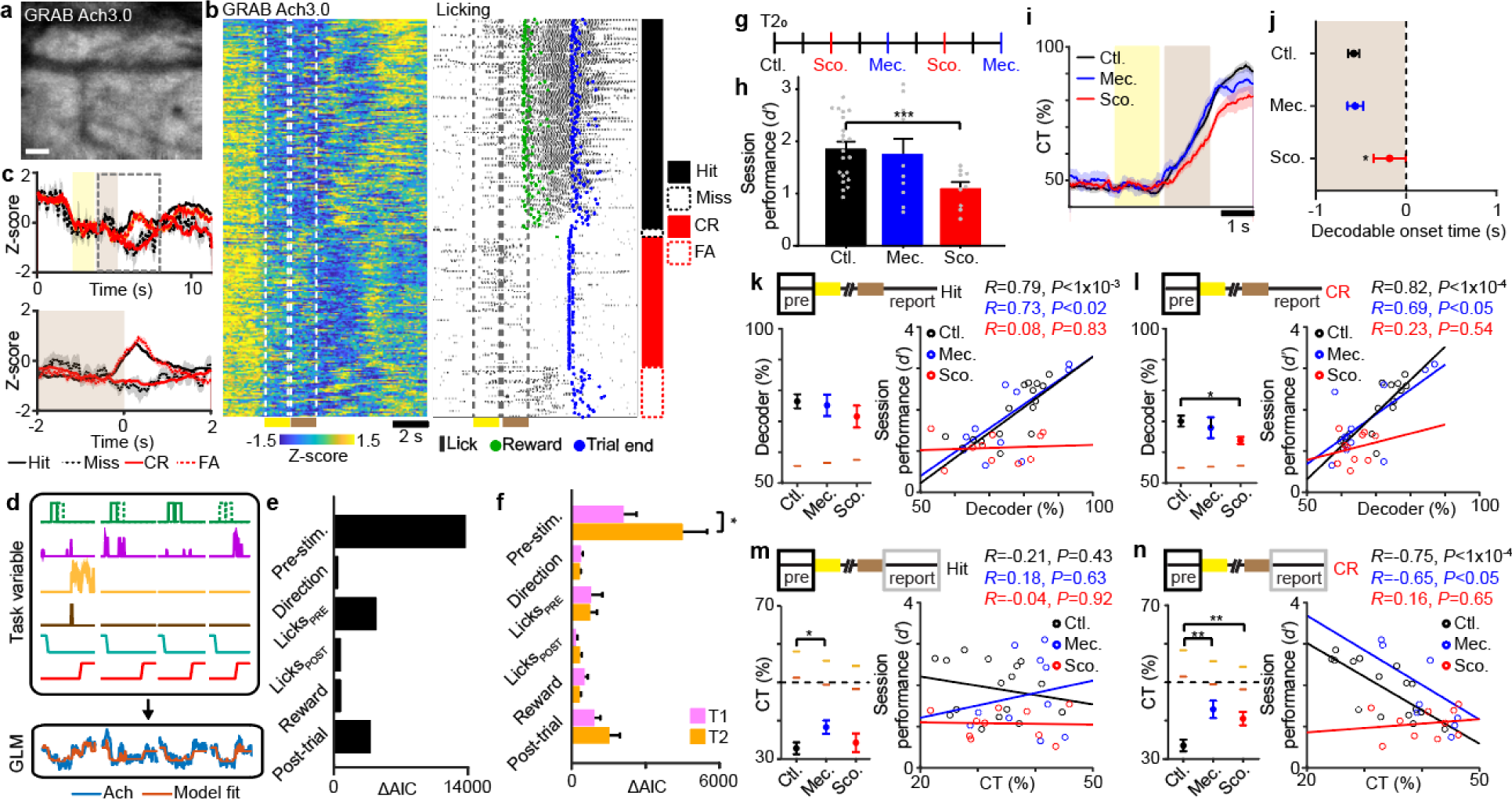
Expected outcome depends on acetylcholine signaling. **a,** Two-photon images of GRAB-Ach3.0 expression in perirhinal cortex. **b,** Example bulk Ach signals (left) and licking behavior (right) sorted by trial type for one session. Timepoint of reward and the end of the trial are also indicated. **c,** Mean Ach signals across the trial period separated by choice aligned beginning of trial (top). Bottom panel shows magnified view of signals (dotted line in top panel) aligned to behavioral report. **d,** Schematic of GLM depicting basis functions for task variables (top) applied to model Ach signals (bottom). **e,** Example encoding of task factors from imaging session shown in [b]. **f,** Encoding of task factors across T1 and T2 sessions. **g,** Schematic of T2 calcium imaging sessions alternating between control no inactivation (Ctl), nAch receptor inactivation by mecamylamine (Mec.), and mAch receptor inactivation by scopolamine (Sco.). **h,** Task performance across pharmacological inactivation sessions. **i,** Stimulus-reward association determined by cross-temporal (CT) decoder performance for Hit vs. non-Hit trials across pharmacological conditions. **j,** Decodable onset timepoint for Stimulus-reward association for [i] across pharmacological conditions. **k,** Decoder performance to Expected Hit (left) across pharmacological conditions. Scatter plot (right) correlation to task performance for individual behavior sessions. **l,** Decoder performance to Expected CR (left) across pharmacological conditions. Scatter plot (right) correlation to task performance for individual behavior sessions. **m,** Cross-temporal (CT) performance to Expected Hit (left) across pharmacological conditions. Scatter plot (right) correlation to task performance for individual behavior sessions. **n,** CT performance to Expected CR (left) across pharmacological conditions. Scatter plot (right) correlation to task performance for individual behavior sessions. Error bars = SEM. Scale bar = 20µm. \**P*<0.05, \*\**P*<0.01, \*\*\**P*<1×10^−4^. *n*=4 animals, 29 T1, 26 T2 sessions for [f]; *n*=4 animals, 19 Ctl., 10 Mec., 11 Sco. sessions for [g-n].

Ach modulates neuronal activity via either nicotinic (nAch) or muscarinic (mAch) receptors^22^. To determine if sustained expected outcome depends on a specific Ach receptor, two-photon calcium imaging was performed on animals trained up through T2. Using reversible pharmacological treatments, population activity was monitored while nAch or mAch receptors were inactivated using mecamylamine or scopolamine, respectively. Inactivation occurred in alternating imaging sessions that were additionally interleaved with control recovery sessions (Fig. 8g). We found that systemic administration of scopolamine, but not mecamylamine, significantly impaired task performance (Fig. 8h, *P*<1×10^−4^, Student’s *t-*test). Population activity was also disrupted. Using a cross-temporal decoder trained on Hit vs. no-hit trials during the report period, we find that scopolamine treatment weakened stimulus-reward associations (Fig. 8i,j). Compared to control conditions, the onset of decodable reward outcome was delayed with scopolamine (*P*<0.02, Student’s *t*-test). No difference was observed with mecamylamine.

We next examined how nACh or mACh receptor inactivation affected expected outcome activity and how those activity patterns related to task performance. Pharmacological blockade did not affect Expected Hit (Fig. 8k). However, while the strength of the decoder correlated with behavioral performance under control and mecamylamine conditions, no significant relationship was observed under scopolamine conditions (*R*=0.08, *P*=0.83, Pearson’s correlation). Scopolamine weakened decoder performance for Expected CR (*P*<0.02, Student’s *t-*test) and its correlation with behavioral performance. (Fig. 8l, *R*=0.23, *P*=0.54, Pearson’s correlation).

To determine whether the sustained property of expected outcome activity was also disrupted, we examined cross-temporal performance for Hit and CR decoders. Mecamylamine weakened the below chance Hit and CR cross-temporal performance (Hit: *P*<0.05, CR: *P*<0.002, Student’s *t-*test). Scopolamine only weakened CR cross-temporal performance (*P*<0.01, Student’s *t-*test). While cross-temporal decoder performance for Hit trials did not correlate with behavioral performance across any conditions (Fig. 8m), CR trial performance was negatively correlated with task performance under control and mecamylamine conditions (Fig. 8n). This was disrupted under scopolamine conditions (*R*=0.16, *P*=0.65, Pearson’s correlation). Overall, this demonstrates that both nAch and mAch receptor-mediated signaling are involved in establishing sustained expected outcome activity in Prh. This expected outcome activity is necessary for correct task performance.

## DISCUSSION

In summary, we demonstrate how Prh is involved in learning an internal model of sensory-guided task behavior that we refer to as a predictive map. Through chronic chemogenetic inactivation of Prh during automated home-cage training, we show that Prh is involved in sensory-guided task learning. While home-cage training with animals under freely moving conditions enable high-throughput, unbiased assays of complex task learning, a limitation of this approach with respect to this study is that the behavioral conditions are not identical to the head-fixed conditions used for characterizing Prh calcium and Ach responses. While experimental differences exist between freely moving and head-fixed tasks, the role of Prh has been demonstrated under other task conditions^16, 18^, reinforcing the idea that Prh supports sensory learning across multiple behaviors. Our analysis of sensorimotor variables during head-fixed conditions along with Prh activity as described below indicates that Prh neurons do not encode sensory and motor information in a direct, bottom-up manner as observed in primary somatosensory cortex^28, 32, 34^. Instead, we propose that sensory information is transformed in Prh into a predictive map that is reflected in three forms of activity: 1) sensory prediction errors; 2) stimulus-outcome associations, and; 3) expected outcome signals (Extended Data Fig. 12).

Sensory prediction errors reflect the learning of task relevant stimulus features. We show that information about stimulus direction - a task relevant feature - decreases with learning but is still present in error trials. Stimulus speed information – corresponding to the strength in stimulus direction - increases with learning and is accompanied by higher firing rates on error trials. These changes with learning are consistent with theories of predictive coding in which neurons signal the difference between expected and actual sensory information^1^. We speculate that Prh evaluates an internal model of task-relevant stimuli via the hippocampus against ongoing stimuli information from sensory neocortex resulting in signals that reflect sensory prediction errors. These results are consistent with previous studies attributing Prh’s role in novel object recognition memory^20, 21^, wherein familiarity is learned from repeated exposure to objects such that novel objects signal the prediction error between experienced and familiar stimuli. In our experimental design, animals experienced slow directions at lower frequencies than fast directions. This does not allow us to disambiguate whether the sensory prediction error signals we observe are driven by familiarity due stimulus probability or task-dependent feature learning. However, our computational model developed for familiarity detection^36^ and applied to recapiluate our experimental results suggests that both phenomena could arise from similar mechanisms.

Sensory prediction errors in Prh may serve two purposes. First, they may act as a teaching signal that promotes updating of task-related variables through error-driven learning that functions to minimize differences between actual and expected sensory information^11^. This would produce a more accurate internal model of task relevant sensory features. Second, considering feedback connections from Prh back to sensory neocortex, prediction errors may aid in sensory inference by boosting bottom-up sensory information in lower areas under circumstances of discrepant sensory signals to help guide behavior^43^. Our results suggest a relationship between the strength of prediction error signals and incorrect choice behavior. Inference may help to support feature invariant encoding of task relevant stimuli (ie. encoding direction invariant to speed).

While stimulus features that are necessary but not sufficient to predict outcome are encoded as sensory prediction errors, combined features that are sufficient to predict reward are encoded as stimulus-reward associations. Through task learning, stimulus-reward associations stabilize and expand in a retrograde manner from the time of reward back to the test period. These signals show similarity to goal-approach neurons in the medial entorhinal cortex and hippocampus during spatial navigation behavior, which increase their activity as animals approach learned locations of reward^44^. This representation generalizes to novel stimulus-reward contingencies. New associations distinctly emerge through a similar mechanism of retrograde expansion. The novel contingency then geometrically aligns with existing associations into an abstract format^7^. This demonstrates that predictive maps can flexibly adapt to newly encountered stimulus-reward contingencies.

Finally, we observe sustained network activity that links prospective signals of expected outcome with the experienced outcome. These signals, along with stimulus-reward associations, depend on cholinergic signaling. More specifically, blockade of mAch receptors disrupts this sustained link in Prh as well as task performance. We speculate that expected outcome signals facilitate learning and recall of sensory-related task models^20, 21, 45^. Ach is released at the beginning of each trial to establish a task-specific expected outcome state space. High cholinergic tone has been associated with an encoding-like “external” mode of processing in the hippocampus and neocortex while low Ach is associated with a retrieval-like “internal” mode of processing^24^. We propose Ach-associated, expected outcome activity may enable sensory information to be evaluated against internal models underlying prediction coding and error-driven learning, consistent with an external mode of processing. Once sensory evidence is sufficient to predict reward, the network switches to retrieval-like “internal” mode in which stimulus-reward associations are retrieved from long-term memories ascribed to cognitive maps. Thus, a predictive map of task behavior could emerge from these switches in network states that engages other brain areas and allows error-driven and associative plasticity to guide model learning in local circuits.

## Acknowledgements

We thank Y. Livneh for guidance in prism implant surgeries, A. Dong, F. Wang, A Deng for assistance in home cage training system design, A. Williams, M.W. Howard for guidance in data analysis, M. Hasselmo for comments on the manuscript. This work was supported by grants from the Richard and Susan Smith Family Foundation (J.L.C.), Elizabeth and Stuart Pratt Career Development Award (J.L.C.), Whitehall Foundation (J.L.C.), Harvard NeuroDiscovery Center (J.L.C.), Boston University Kilachand Fund Award (J.L.C.), National Institutes of Health BRAIN Initiative Award R01NS109965 (J.L.C.), National Institutes of Health New Innovator Award DP2NS111134 (J.L.C.). NSF Neuronex 1707398 (S.F.); the Gatsby Charitable Foundation GAT3708 (S.F.), the Simons Foundation (S.F.), the Swartz Foundation (S.F.). D.G.L., C.A.M., and J.L.C. designed the study. D.G.L. and G.H. designed the home cage training system, D.G.L., A.E.C., and G.H. performed home cage inactivation experiments, D.G.L., C.A.M., and O.K. performed two-photon imaging experiments. R.N. performed computational modeling. G.D.L. performed retrograde tracing experiments. D.G.L., D.L.M, and J.L.C. performed data analysis. R.N., S.F., and J.L.C. supervised data analysis. D.G.L., C.A.M., and J.L.C. wrote the paper.

## METHODS

### Mice

Experiments in this study were approved by the Institutional Animal Care and Use Committee at Boston University and conform to NIH guidelines. Behavior experiments were performed using male and female C57BL/6J mice (The Jackson Laboratory). All animals were 6-8 weeks of age at time of surgery. Mice used for behavior were housed individually in reverse 12-hour light cycle conditions. All handling and behavior occurred under simulated night time conditions.

### Animal preparation

Prh was targeted stereotaxically (2.7 mm posterior to bregma, 4.2 mm lateral, and 3.8mm ventral). For inactivation experiments, bilateral injections were targeted via the parietal bone. For each side, animals received either retroAAV-*hSyn*-*Cre* (4.5×10^12^ vg/mL) and AAV9-*hSyn*-*dio-hM4Di-mCherry* (6.0×10^12^ vg/mL) (1:1, 600nL) or retroAAV-*hSyn*-*Cre* and AAV9*-hSyn-dio-mCherry* (6.0×10^12^ vg/mL) (1:1, 600nL). For tracking in the home cage training, a radio frequency identification (RFID) glass capsule (SEN-09416, Sparkfun) was implanted subcutaneously in the animal’s back. For *in vivo* imaging experiments, a unilateral injection was targeted via the temporal bone at 250 µm and 500 µm below the pial surface of either AAV.PHP.eB-*EF1*α*-RCaMP1.07* (600nL, 6×10^12^ vg/mL), AAV9-*hSyn-GRAB-Ach3.0* (600 nL, 2.5×10^12^ vg/mL), or AAV2-retro-*CAG-GFP* (600nL, 1×10^12^ vg/mL). For optical access, an assembly consisting was of a 2 mm aluminum-coated microprism (MPCH-2.0, Tower Optical) adhered to coverglass along the hypotenuse and the side facing Prh was implanted over the pial surface. A metal headpost was implanted on the parietal bone of the skull to allow for head fixation. For unilateral retrograde tracing between Prh and S2, CTB-Alexa647 (Molecular Probes, Invitrogen; 300 nL, 1% wt/vol) was delivered into Prh, targeted via the temporal bone and CTB-Alexa488 (300 nL, 1% wt/vol) was delivered into S2 (0.7 mm posterior to bregma, 4.2 mm lateral, 250 and 500 µm below the pial surface).

### Home cage task training

Two weeks after injections, animals were trained to a whisker-based context-dependent sensory task adapted for training in an automated live-in environment (**Supplementary Text S1**). The animals were singly housed in individual cages. Three cages were attached to a shared training system wherein individual access was restricted via servo-operated doors (SG92R, Tower Pro) controlled by a microcontroller (Uno Rev3, Arduino). The training system consists of a narrow corridor that restricts body and head movement at the front of the corridor where sensory stimulus is delivered. Equipment for whisker stimulus, lick detection, sound delivery, air puff delivery, and water delivery were similar to as described^28^. Water ports were attached to a capacitive lick sensor (AT42QT1010; SparkFun) that dispenses 5 to 6 uL of water through a miniature solenoid valve (LHDA0531115H; The Lee Company). For the rotation stimulus, commercial grade sandpaper (3M; roughness: P100) was mounted along the outside edge of a 6 cm diameter rotor, attached to a stepper motor (Zaber) to deflect the whiskers which was mounted onto a linear stage (Zaber) to place the rotor within whisker reach. Two lick ports were mounted onto a linear actuator (L12-P, Actuonix) that controlled access to water during the task. An LED beam breaker (2167, Adafruit) at the head of the training system such that animals self-initiated behavioral trials by breaking the beam with their body.

Each animal was provided access to the training system via the servo door through scheduled two-hour morning and two-hour afternoon session blocks. Animals were initially acclimated by learning to retrieve water from the lick ports. Once acclimated, animals proceeded to task training. During task training, the rotor providing whisker stimulus was retracted during the inter-trial interval and placed in reach during stimulus periods. The lick spouts were only presented during the report period and retracted at all other times. A two-forced alternative choice task design was used in which correct choice required licking to the right port for non-match stimuli and to the left port for match stimuli. Only fast rotations (1.75 cm/s) of stimulus direction were used.

Training was divided into 5 stages (T1-T5) (Table 1, **Supplementary Text S3**). For T1 and T2, one non-match stimuli (AP) and two match stimuli (AA, PP) were included. T1 was defined as initial naïve performance. T2 was defined as learned performance beginning from the point in which animals displayed *d’* > 0.45 for two consecutive sessions. For T3, the second non-match stimuli (PA) was introduced. For T4, delays between the sample and test stimuli were gradually lengthened up to 2 seconds. The rotor was also gradually retracted up to 1.5cm out of whisker reach. T5 was defined as consistent expert performance with 2s delay and 1.5cm rotor retraction. Advancement from T2-T5 was automated based on behavioral performance of two consecutive sessions of >80% correct (*d’* ∼1.68). The delay period and rotor withdrawal distance during T4 was automatically increased based on behavioral performance of >80% correct (*d’* ∼1.68) across a 15-trial sliding window.

In addition to water reward, correct behavioral choice was reinforced using three automatically adjusted task settings (Table 3, **Supplementary Text S4**). Punishment in the form of a combination of time outs (2-10s) and air puffs to the face were introduced to discourage incorrect decisions. Time outs ranged from 2-10s. Air puffs (100ms) ranged from 1-5 trains and were introduced for >7s time out. Punishment systematically increased during poor performance corresponding to <70% correct (*d’* ∼1.05) over a 50 trial sliding window. Punishment was automatically decreased if the proportion of misses in this window exceeded 50%. To correct for report biases in which animal repetitively licked one port irrespective of stimulus condition, the probability of match vs. non-match stimulus conditions was increased in favor of the stimulus condition associated with the neglected spout. To correct for primacy and recency stimulus bias resulting in disproportionally greater error trials for one of the two match conditions or one of the two non-match conditions, probability of one of the two match or non-match conditions was adjusted in favor of the condition with the greater proportion of errors.

For chemogenetic inactivation, Compound 21 (HB6124, HelloBio) was provided in the drinking water (9.5µg/mL H_2_O, 1mg/kg body weight). Animals only received water by performing the task. Their weight was monitored daily to ensure body weight did not drop below 80% of initial weight. Animals were trained continuously for six weeks.

### Head-fixed task training

Two weeks after microprism implantation and injections, animals were handled and acclimated to head fixation. Training to a head-fixed whisker-based context-dependent sensory task was performed similar to as described^28^ (**Supplementary Text S2**). Water ports and stimulus delivery hardware were same as the home-cage training system. Whiskers were trimmed to a single row for videography. Animals trained for two sessions per day. A go/no-go task design was used in which animals licked for water reward for non-match stimulus conditions and withheld licking for match stimulus conditions. T1-T3 training stages were similar as stages defined in home cage task training (Table 2). For T4, the delay between sample and test stimuli was gradually increased from 100ms to 2s with the rotor remaining within whisker reach through the delay period. For T5, the rotor was retracted 1.5cm during the delay period across delays of 2s, 3s, and 4s which were randomly presented with probabilities of 50%, 25%, and 25% respectively. Fast (1.75 cm/s) and slow (0.87 cm/s) rotations of stimulus direction were used. For T1-T4, slow directions represented 5% of all trials. For T5, the fraction of slow trials was increased to 25% of all trials

Adjustments to task settings to reinforce correct behavioral choice were carried out semi-automatically. Punishment in the form of a combination of time outs (2-10s) and air puffs (100ms) ranging from 1-5 trains to the face was manually adjusted to discourage false alarm licking on match trials. During T1, the probability of non-match stimulus conditions was manually reduced to 35-40% of all trials reduce false alarm trials or increased up to 60% to reduce miss trials. To correct for primacy and recency stimulus bias resulting in disproportionally greater error trials for one of the two match conditions or one of the two non-match conditions, probability of one of the two match or non-match conditions was adjusted in favor of the condition with the greater proportion of errors. Animals only received water by performing the task. Their weight was monitored daily to ensure body weight did not drop below 80% of initial weight. Animals were trained continuously and terminated once animals had performed at least 4-6 T5 sessions.

### Acetylcholine receptor inactivation

Microprism implanted animals expressing RCamp1.07 in Prh were imaged and trained up to expert T2 performance. Mecamylamine (1mg/kg b.w.) or scopolamine (1-5 mg/kg b.w.) was delivered systemically vis intraperitoneal (IP) injection ∼1h prior to behavior imaging session. For control conditions, behavior imaging sessions was performed at least 16 hours after the previous pharmacological inactvation session to allow for recovery.

### Histology

Mice were anaesthetized (sodium pentobarbital; 100 mg per kg and 20 mg per kg body weight) and perfused transcardially with 4% paraformaldehyde in phosphate buffer, pH 7.4. For anatomical tracing experiments, coronal sections (50-75 µm) were cut using a vibratome (VT1000; Leica). For chemogenetic inactivation experiments, coronal sections (150 µm) were cut, tissue cleared and embedded in hydrogel using PACT-CLARITY, and stained for *Fos* (B4-Alexa647 hairpin amplifiers) using HCR-FISH as previously described^27^. Images were acquired using a epifluorescent microscope (Eclipse NiE, Nikon) or spinning disk confocal microscope (Ti2-E Yokogawa Spinning Disk, Nikon).

### Two-photon imaging

Two-photon calcium imaging was performed with a custom-built resonant-scanning multi-area two-photon microscope with a 10x/0.5NA, 7.77mm WD air objective (TL10X-2P, Thorlabs) using custom-written Scope software^33^. A 31.25 MHz 1040 nm fiber laser (Spark Lasers) was used for RCaMP1.07 imaging. Simultaneous imaging at 32.6 Hz frame rate was performed of two imaging planes in L2/3 separated 50 µm in depth. For GRAB-Ach3.0 or GFP imaging, a single area at 32.6 Hz frame rate was acquired using an 80MHz ti:sapphire laser (Mai Tai HP DeepSee, Spectra Physics) tuned to 950 nm. Average power of each beam at the sample was 50-90mW. Imaging was performed during head-fixed task behavior or during passive stimulation sessions in naïve animals using similar stimulus conditions as T5.

### *In vivo* image analysis

All image processing was performed in MATLAB, Python, and ImageJ as described^28, 46^. For calcium imaging analysis, two-photon images were first motion corrected using a piece-wise rigid motion correction algorithm^47^. Independent noise related to photon shot noise was removed from the image times-series using DeepInterpolation^48^. To identify neurons chronically imaged across all behavior sessions, a global reference image was generated by tiling FOV images from each session to account for slight variations in positioning and to reveal a common FOV shared by all sessions. ROIs were manually identified by comparing structural images based on fluorescence intensity and a map of active neurons identified by constrained non-negative matrix factorization from image time series. ROI positions were adjusted for each session to account for tissue changes or rotations over longer time scales. Calcium signals were then extracted for each ROI for each session. A global neuropil correction was performed for each neuron and the resulting fluorescence traces were detrended on a per trial basis. For acetylcholine imaging analysis, the fluorescence intensity across the entire FOV was averaged to obtain a bulk signal of Ach dynamics. Ach signals were z-scored on a per trial basis.

### Calcium event estimation

Calcium signals were deconvolved using an Online Active Set method to Infer Spikes (OASIS), a generalization of the pool adjacent violators algorithm (PAVA) for isotonic regression^49^. First, calcium signals below baseline fluorescence (bottom 10^th^ percentile of signal intensity) were thresholded. For each cell, a convolution kernel with exponential rise and decay time constants were determined using an autoregressive model. For measurement of photon shot noise, signal-to-noise (*v)* was calculated as for each cell:

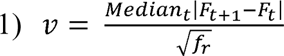

where the median absolute difference between two subsequent time points of the fluorescence trace, *F*, is divided by the square root of the frame rate, *f* ^50^. The convolution kernel was applied to the calcium signals to obtain an initial deconvolved signal that was then normalized by the signal-to-noise resulting in a calcium event estimate (*ŝ*).

### Population decoding analysis

To decode population activity with respect to trial conditions, maximum margin support-vector machine (SVM) linear classifiers were used on the single-trial population response vectors of simultaneously recorded neurons within one imaging session^7^. For each neuron in the population, calcium events across a given time window was averaged for each trial and then z-scored across all trials in session time. For each classifier, activity from 10-20% of trials was separated for testing while the remaining trials were used to train the classifier. In the case of comparing stimulus direction or reward, in which >100 trials were recorded for each condition (i.e., anterior vs. posterior for stimulus direction or hit vs. non-hit), the accuracy of the decoder performance was determined using 10-fold cross validation. For comparing stimulus speed or choice in which slow speed conditions or error conditions were very few or varied across task learning (Fig. 3, Extended Data Fig. 9), trials in the minority condition in the training set were randomly resampled to match trial numbers in the other condition before 10-fold cross validation. This process was repeated 100 times and the decoder accuracy was calculated from the average accuracy. The statistical significance of the decoding accuracy was assessed by shuffling the trial labels in the training set prior to classification. This process was repeated 1000 times and decoder accuracies above the 95^th^ or below the 5^th^ percentile of the shuffled distribution was determined to be statistically significant.

For a cross-temporal classifier (Figs. 5-8), SVMs were trained as described above using average activity across the pre-stimulus period, sample period, test period, report period, or a sliding window of 1000 milliseconds. The cross-temporal accuracy was determined using 10-fold cross-validation by testing on withheld trials from activity across different pre-stimulus period, sample period, test period, report period, or a sliding window of 300 milliseconds. Significant cross-temporal decoding was determined by shuffling the population vector weights and then testing performance on the resulting shuffled decoder. This process was repeated 1000 times and cross-temporal accuracies above the 95^th^ or below the 5^th^ percentile of the shuffled distribution was determined to be statistically significant. The decodable onset of the reward outcome classifier was defined as the first significant timepoint across the test and report period.

For a cross-session classifier (Fig. 5), SVMs were trained using average activity across the pre-stimulus or report period consisting of 80-90% trials from one imaging session. The cross-session accuracy was determined using 10-fold cross-validation by testing on average activity in the same trial period window in a different session using all trials. The same neuronal population imaged across sessions was used for training and testing. Significant cross-session decoding was determined by shuffling the population vector weights and then testing performance on the resulting shuffled decoder. This process was repeated 1000 times and cross-session temporal accuracies above the 95^th^ percentile of the shuffled distribution were determined as statistically significant.

For cross-condition analysis of rewarded stimulus conditions (Fig. 6), non-match stimulus trials were separated by stimulus condition (anterior-posterior or posterior-anterior) into a training or testing set. Match stimulus trials were randomly separated into the training or testing set. SVMs were then trained using average activity from the report period along hit vs. non-hit trial conditions. The cross-temporal accuracy of the cross condition was determined using 10-fold cross-validation by using the average activity across a sliding window of 300 milliseconds of the test set. The cross-temporal accuracy at 300ms from the end of the test period was used to assess the strength of the cross-condition of the test period.

### Choice selectivity

To determine the relationship between stimulus speed encoding and choice selectivity, an SVM was trained to speed trials. Neurons with significant population vector weights were determined by shuffling the trial labels in the training set prior to classification. This process was repeated 1000 times to obtain a shuffled distribution for each neuronal weight. Neuron weights above the 95^th^ or below the 5^th^ percentile of the shuffled distribution were determined to be statistically significant. For significant neurons, selectivity to correct (hit, correct rejection) or error (miss, false alarm) trials was determined by calculating the average event rate for each of the two trial conditions. The peak activity level during either the sample or test period as measure of a neuron’s stimulus response (SR). Choice selectivity was expressed as *(SR_ERROR_ - SR_CORRECT_)/(SR_ERROR_ + SR_CORRECT_)* where *SR_ERROR_*is the peak response on error trials and *SR_CORRECT_* is the peak response on correct trials.

### Computational modeling

An autoencoder was trained to reconstruct a two-dimensional input signal (Fig. 4). The input signal consisted of two independent variables, direction of movement and speed, with two different values each. This made a total of four experimental conditions: anterior direction and low speed, posterior direction and low speed, anterior direction and fast speed, and posterior direction and fast speed. These four experimental conditions were mapped to four points on a two-dimensional space [-1,-1], [-1,1], [1,-1], [1,1]. Simulations of *k* trials per experimental condition were performed, producing a total of 4*k* trials (*k* = 100). On each trial additive Gaussian noise with mean zero and variance σ^2^_inp_ was added to the experimental conditions and then expanded by a random projection to an N_inp_ space (σ^2^_inp_ = 0.5, N_inp_ = 10).

The autoencoder consisted of input, intermediate, and output layers. Intermediate neurons were ReLU units with noise (additive Gaussian noise, σ^2^_inp_ = 0.5). Additionally, an additional read-out unit was included that read the intermediate layer to classify direction of motion on a trial-by-trial basis. This additional read-out neuron was added to impose an asymmetry between direction of motion and speed in both the intermediate and output layers. The loss function that was minimized through learning was:

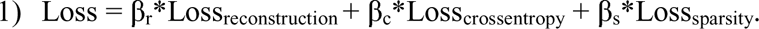

The reconstruction loss was the mean squared error (MSE) between the input and the output layer (β_r_ = 0.0001). The cross-entropy loss corresponded to the classification loss of the additional read-out unit that classified direction of motion from the activity of the intermediate layer (β_c_= 1). Finally, we also added an L2-norm sparsity loss on the activity of the intermediate layer to constrain the activity of the intermediate layer (β_s_ = 10). The autoencoder was trained with stochastic gradient descent (ADAM, *lr* = 0.01, batch size = 10) for 200 epochs. A final downstream unit (logistic regression, sci-kit learn) was added that read out from the familiarity population, that is, the difference between the reconstructed output and the input^36^. An independent classifier was trained on each training epoch. The reported decoding performance on both direction and speed corresponds to the mean across cross-validation iterations (5-fold CV) and independent simulations (n = 50).

Alternative models were trained and analyzed. This includes models containing only reconstruction loss (β_r_ = 1, Extended Data Fig. 8a), reconstruction and cross-entropy with respect to direction (β_r_= 0.0001, β_c_ = 1, Extended Data Fig. 8b), and reconstruction, cross-entropy, and L2 sparsity on the hidden layer (β_r_ = 0.0001, β_c_ = 1, β_s_ = 10, and β_s_ = 100, Extended Data Fig. 8c,d). Modeling was performed in Python and PyTorch. Code is available at github.com/ramonnogueira/AutoPerirhinal.

### Acetylcholine signal analysis

To understand the effects of task-relevant variables on the acetylcholine (Ach) dynamics, we fit a Normal GLM to the normalized Grab-Ach3.0 fluorescence acquired on each trial within a recording session. The model calculates an estimated signal, *ŷ_t_*, using:

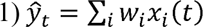

where *x_i_(t)* represents the time course for the *i*^th^ explanatory variable, and *w_i_* represents the weight assigned to this variable relating its estimated effect on the signal^51^. All GLMs were fit using MATLAB’s lassoglm function with a normal distribution, identity link function, 6 penalty values (γ), and 4 fold cross-validation.

Task variables *x_i_(t)* were represented as boxcars corresponding to their occurrence during the time course of a trial. These boxcars had value “true/1” during appropriate time points and “false/0” otherwise. These include “pre-stimulus,” “stimulus direction anterior,” “stimulus direction posterior,” and “post-trial” variables. “Reward” was represented as a boxcar lasting 300ms after the point of reward delivery. Licking events were resampled to match the image acquisition rate. This was then convolved with a 10-sample Gaussian kernel and separated into “pre-reward licking” (Lick_PRE_) and “post-Reward licking” (Lick_POST_) variables based on rewarded trials. All licking on miss, false alarm, and correct rejection trials were considered Lick_PRE_. For hit trials, licks before water reward were Lick_PRE_ while licks after water reward were Lick_POST_.

Related covariates were grouped together into ‘task factors.’ Each task variable was treated as its own “task factor” with the exception of “stimulus direction anterior” and “stimulus direction posterior” which were grouped into a task factor for “stimulus direction.” For each task factor, a partial model was constructed that excluded the covariates associated with this task factor. Any increase in deviance from the full model to the partial model therefore resulted from the exclusion of this task factor’s covariates. Akaike Information Criterion (AIC) was used to compare deviance between partial models in which different number of covariates were excluded such that:

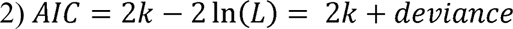

where *k* is the number of model parameters, deviance = *-2*ln*(L),* and *L* is the model likelihood. The difference in AIC (ΔAIC) between the full and partial model was calculated as:

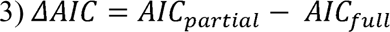

### Statistical procedures

No statistical methods were used to predetermine sample size. For Prh inactivation experiments, investigators were blinded to hM4Di+ or hM4Di-groups during experiments and outcome assessment. For two-photon experiments, animals were not randomized and the investigators were not blinded to allocation during experiments and outcome assessment. Statistical tests used are indicated in figure legends. Error bars on plots indicate standard error of the mean (SEM) unless otherwise noted.

For Prh inactivation experiments, a bootstrap analysis was used to compare the fraction hM4Di+ versus hM4Di-animals able to successfully accomplish the T2 stage. For testing of sequence reliability or stimulus similarity across passive and training stages, a one-way ANOVA was performed followed by a multiple comparisons test. For testing of differences in linear decoder or cross-temporal decoder performance in individual sessions between training stages, a one-way ANOVA was performed followed by a multiple comparisons test. For performance of linear decoders for direction or speed, a Student’s *t*-test was used to compare correct versus error trials at specific training stages. For comparisons of choice selectivity in individual neurons across training stages, a one-way ANOVA was performed followed by a multiple comparisons test. For statistical tests of Ach signal encoding, a repeated-measures ANOVA was performed followed by a multiple comparisons test was used to compare the strength of GLM ΔAIC values between task factors. A Student’s *t*-test was used to compare AP versus PA decoder performance as well as cross-conditional decoder performance at specific T3 sessions. The Bonferroni-Holm method was used to correct for multiple comparisons.

---

**Extended Data Figure 1.**
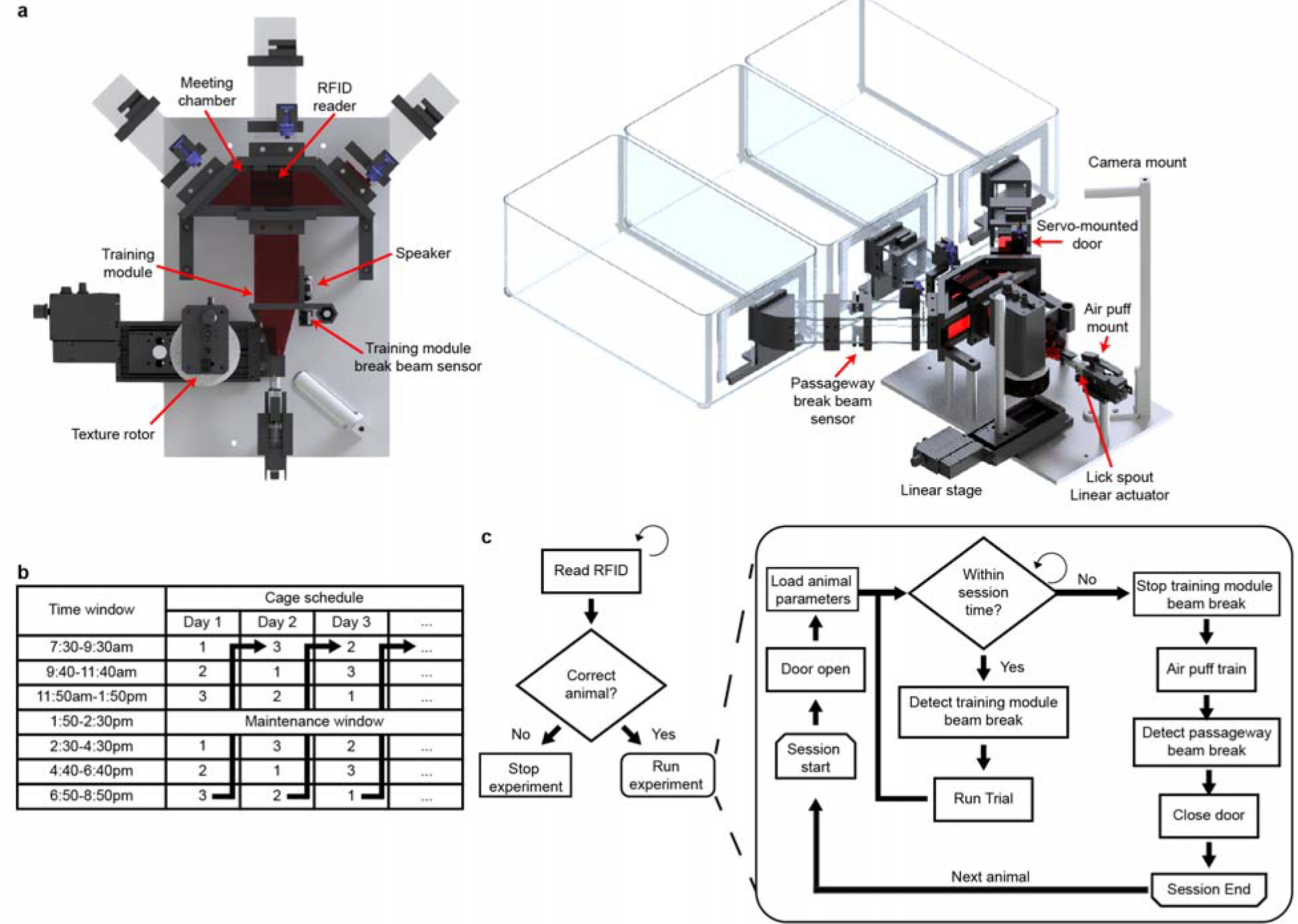
Automated home cage training system. **a,** Mechanical design of home-cage training system designed to support automated training of three individually housed mice. **b,** Rotating daily timetable used for training three animals (1, 2, 3). **c,** Flow chart for managing individual animals in the training system.

**Extended Data Figure 2.**
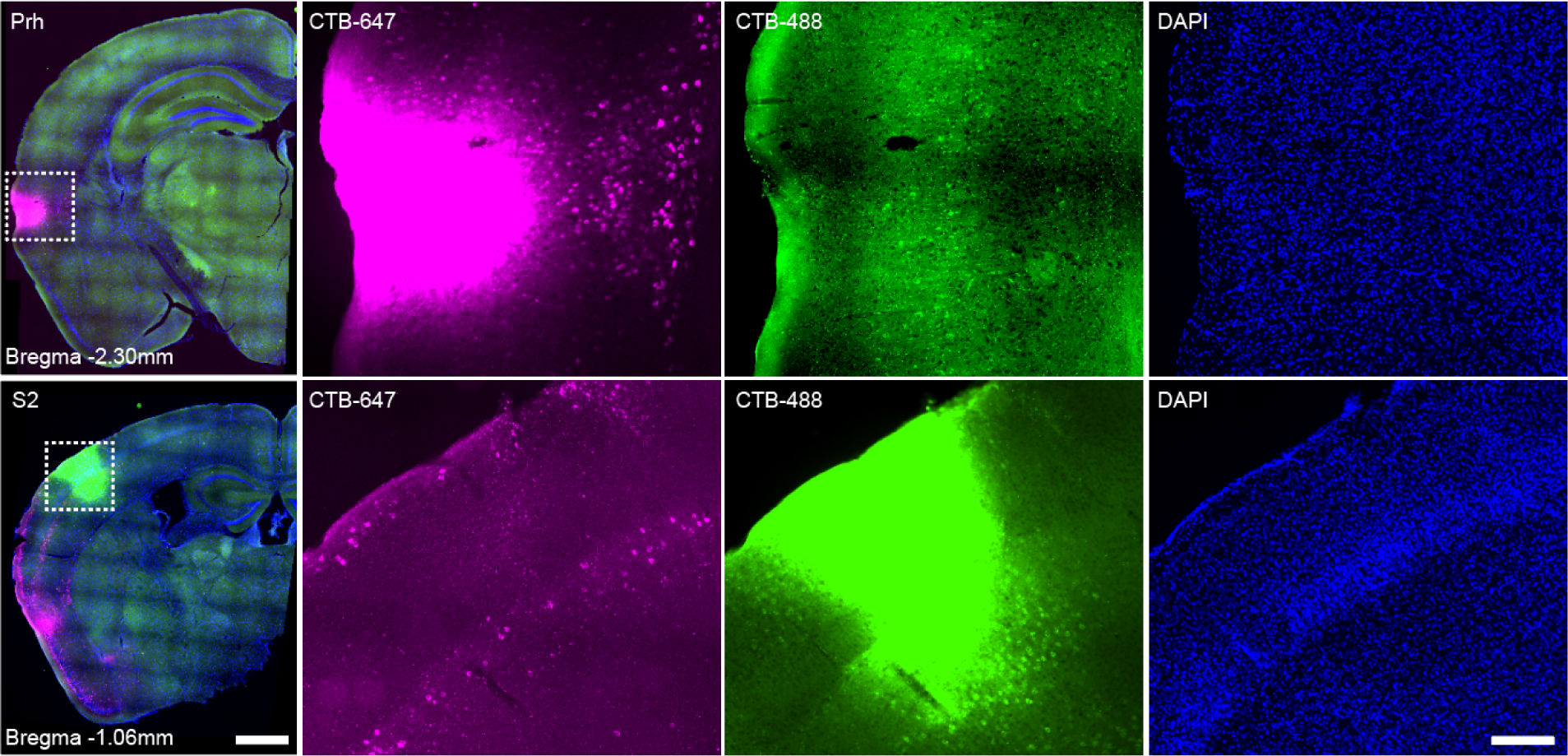
Reciprocal connections between perirhinal cortex and secondary somatosensory cortex. Fluorescent micrographs of coronal sections showing retrograde labeling of projection neurons between perirhinal cortex (CTB-647) and secondary somatosensory cortex (CTB-488). Right panels show magnified view of indicated area in left panel (dotted rectangle). Scale bars: 1mm (left panels), 0.2mm (right panels).

**Extended Data Figure 3.**
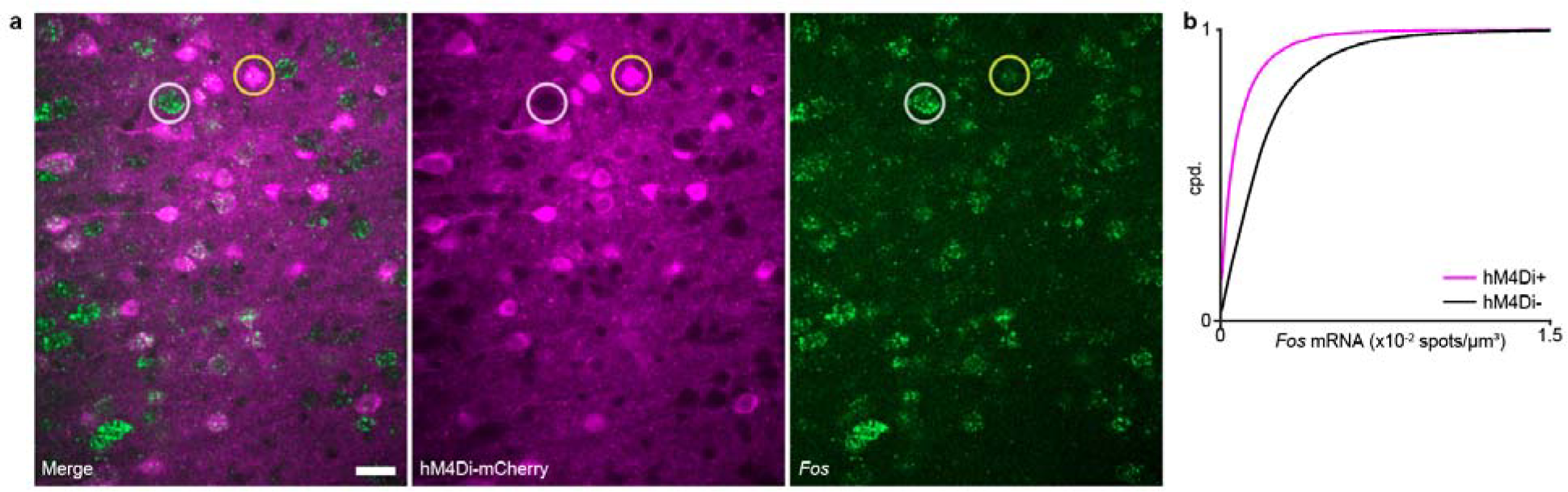
Chemogenetic inactivation of perirhinal cortex. **a,** Validation of chronic inactivation of perirhinal cortex by *Fos* mRNA expression. Animals received Compound 21 in drinking water for up to 6 weeks. *Fos* mRNA was visualized using HCR-FISH. Examples of hM4Di-mCherry+ neurons (yellow) with low *Fos* expression versus hM4Di-mCherry-neurons (grey) with high *Fos* expression are shown. **b,** Cumulative distribution of *Fos* expression measured by HCR-FISH in hM4Di-mCherry+ vs. hM4Di-mCherry-neurons. 74.9±3.0% of neurons were hM4Di-mCherry+, *n* = 4 animals. Scale bar: 20 µm.

**Extended Data Figure 4.**
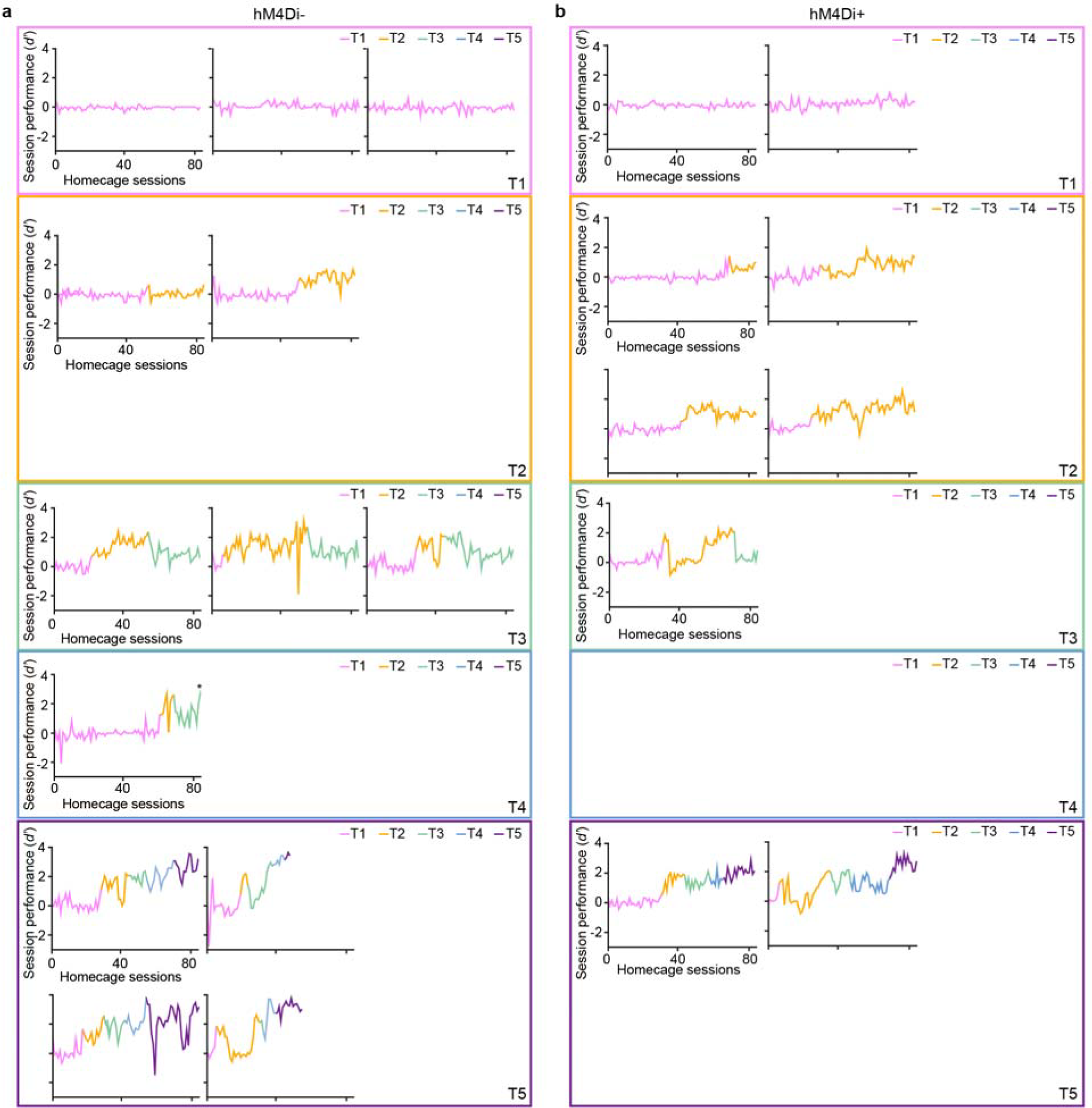
Performance curves for individuals in home cage training task. **a,** Session performance across training for hM4Di-animals sorted by final training stage reached after 84 sessions. Training was stopped prior to 84 sessions for some animals that reached T5. The majority of hM4Di-animals passed T2. The noted animal (*) reached T4 at session 84. **b,** Session performance across training for hM4Di+ animals sorted by final training stage reached after 84 sessions. The majority of hM4Di+ animals failed to passed T2.

**Extended Data Figure 5.**
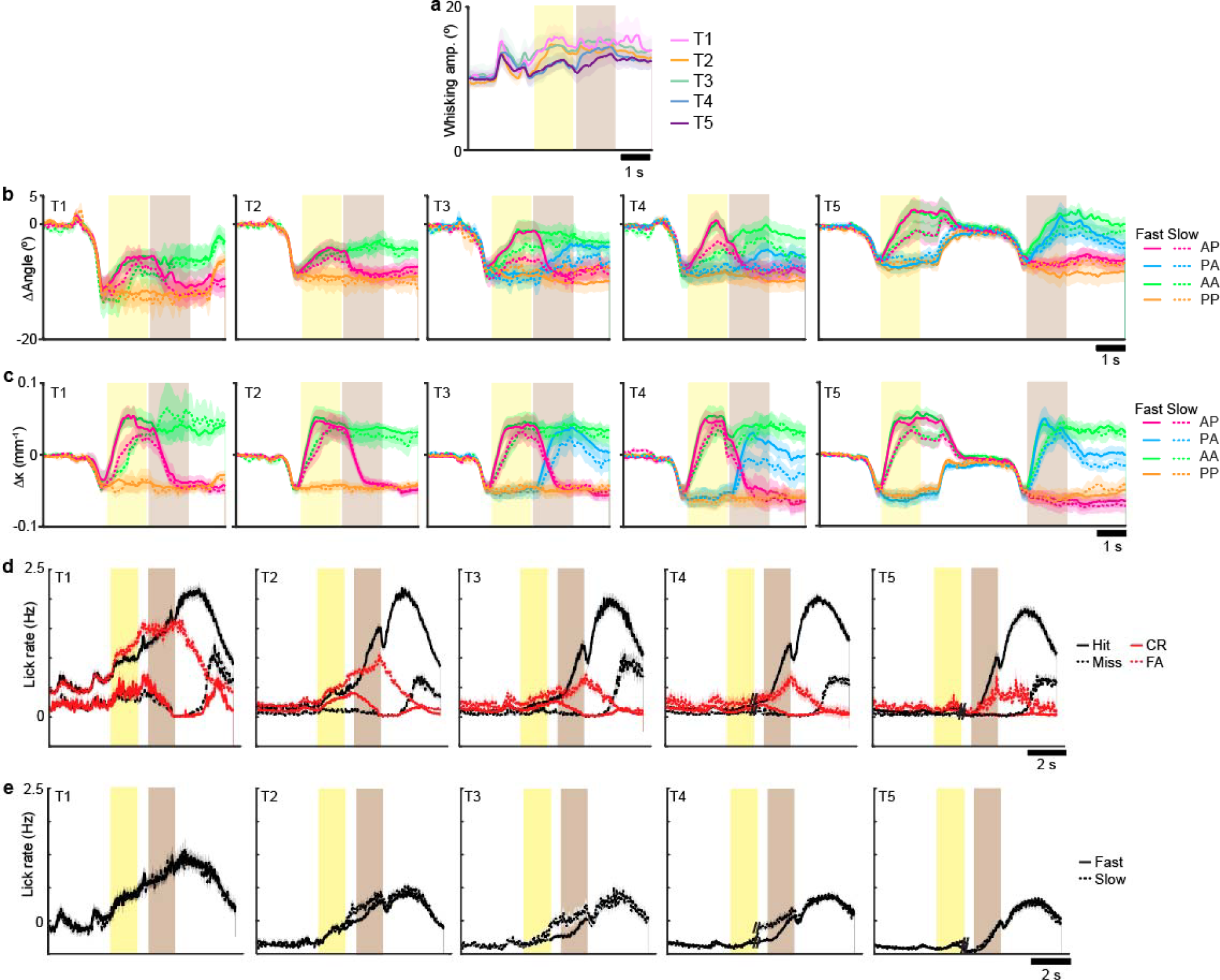
Sensory and motor variables throughout learning. **a,** Mean whisking amplitude over the trial period averaged across training stages. **b-c,** Mean change in whisker angle [b] and curvature [c] by sorted stimulus condition across training stages. **d-e,** Mean lick rate through the trial period across training stages sorted by choice [d] or stimulus speed [e]. Shaded regions = SEM.

**Extended Data Figure 6.**
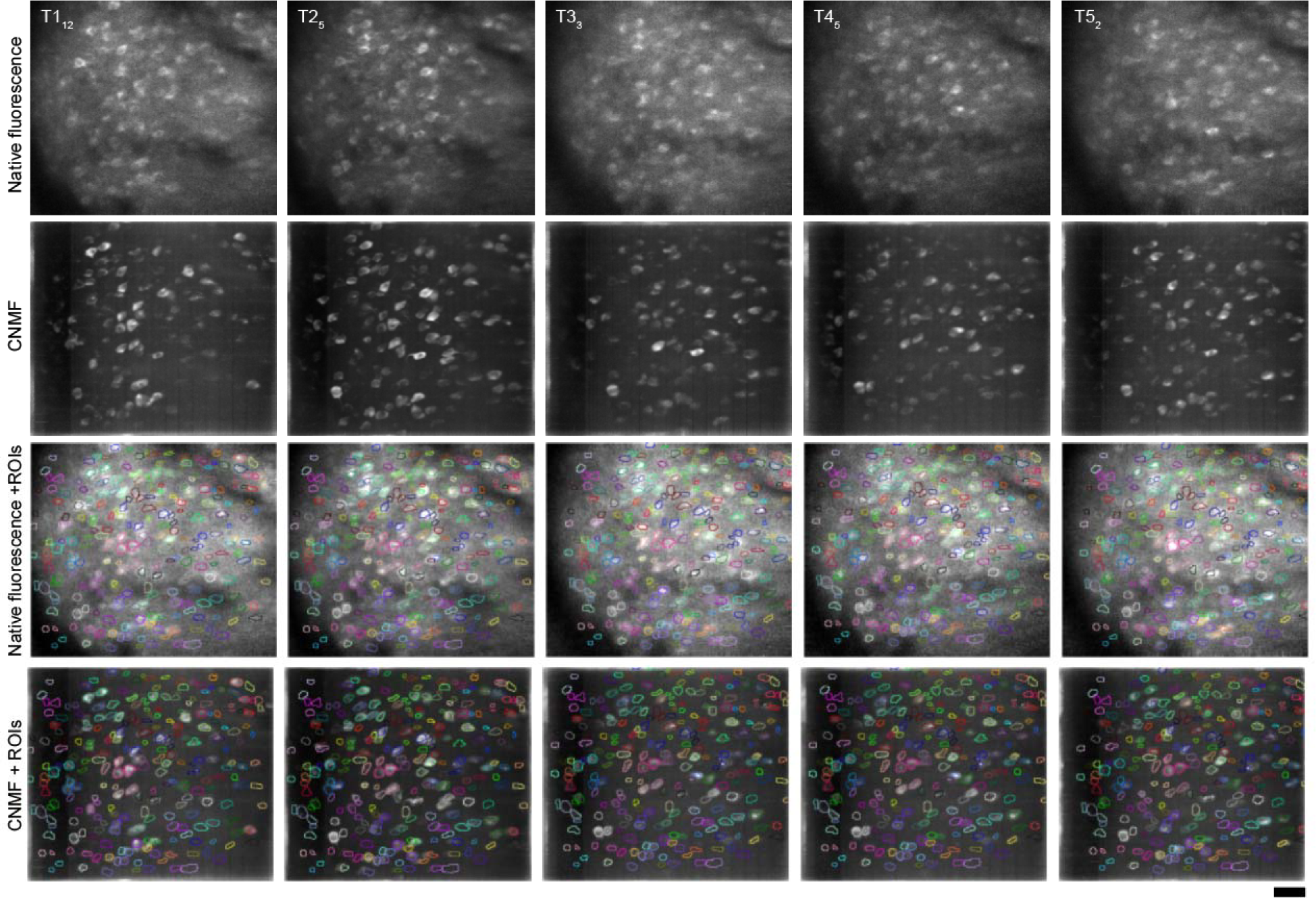
ROI selection across imaging sessions. Examples of ROIs identified throughout the time course of training. ROIs were manually identified and segmented by comparing structural images of native RCaMP1.07 fluorescence and images of ‘active’ neurons through constrained non-negative matrix factorization (CNMF) of the image timeseries across the training session. Structural images were used to identify all neurons (active and inactive) in the session while the CNMF images helped to define boundaries of ROIs. Scale bar: 50µm.

**Extended Data Figure 7.**
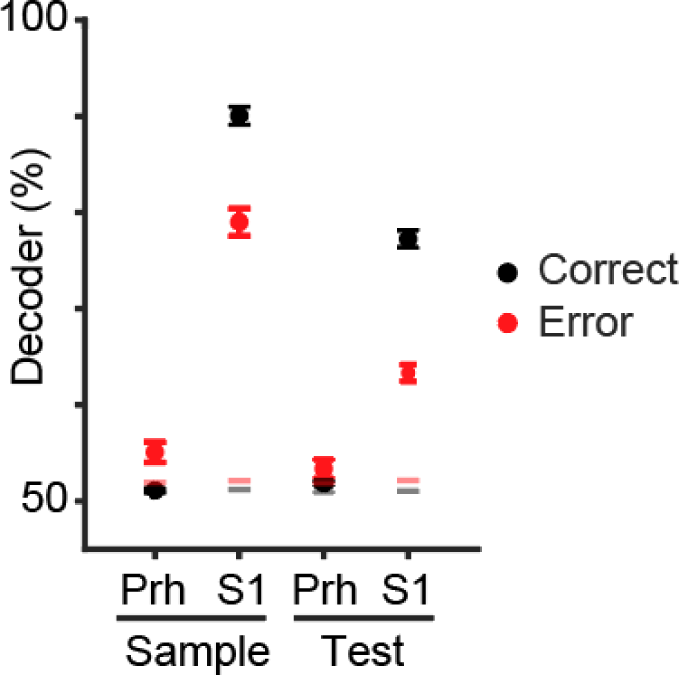
Population encoding of stimulus direction Prh versus S1 in expert animals. Decoder performance on stimulus direction during Sample or Test periods using activity T5 sessions from Prh or S1. S1 neural data was obtained from (ref. 28). Separate decoders were trained and tested using Correct (Hit and Correct Rejection) or Error (False Alarm and Miss) trials. Error bars = SEM. Red and gray bars = 95^th^ percentile of shuffled distribution on Error and Correct trials, respectively.

**Extended Data Figure 8.**
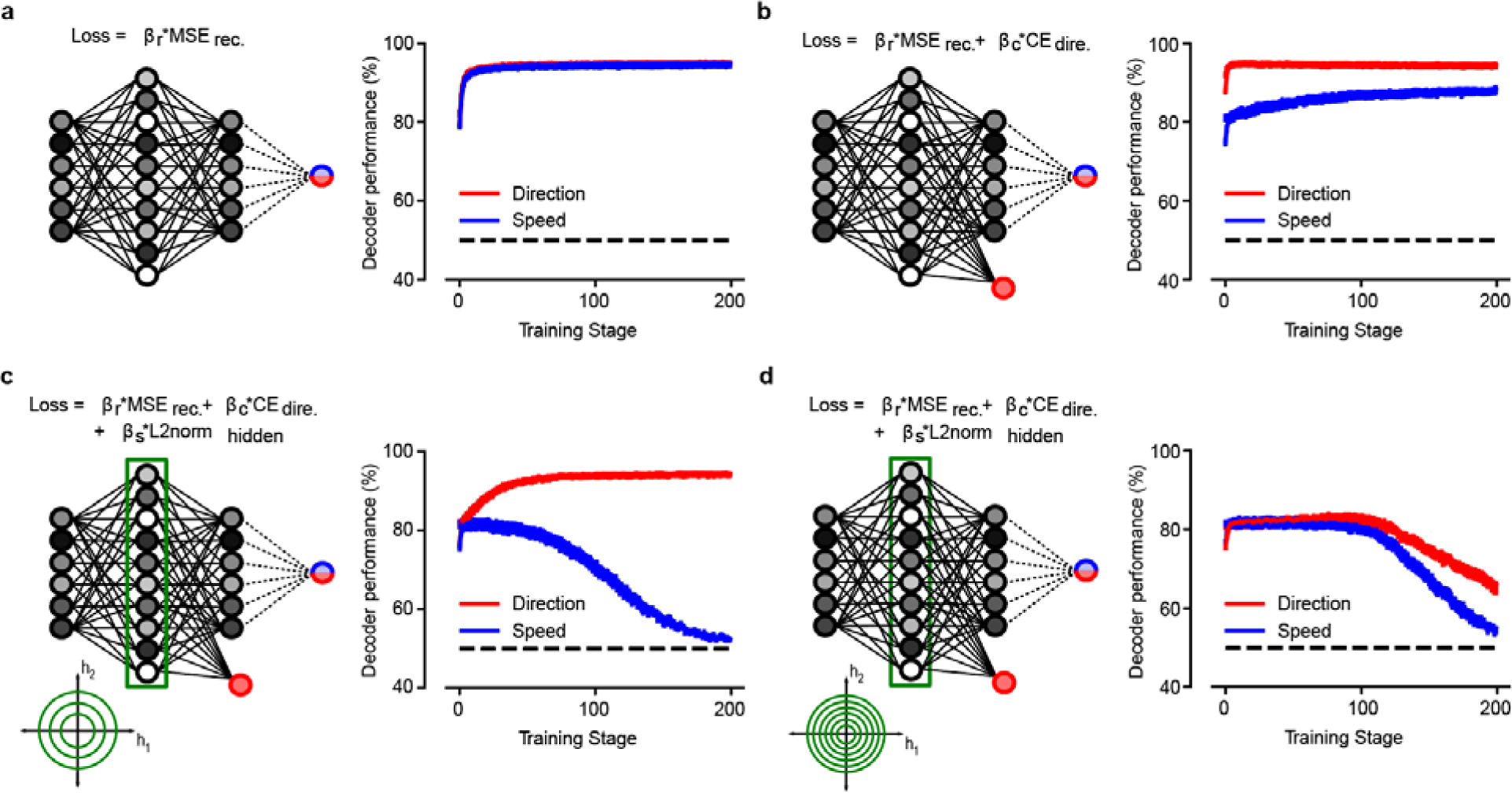
Alternate models produce different decoding performance of direction and speed across learning. Decoding performance for direction of motion (red) and speed (blue) of a downstream neuron that reads out the output layer of the autoencoder (logistic regression, sci-kit learn). **a,** Results from an autoencoder trained to minimize only reconstruction loss (Mean Squared Error, MSE). Direction and speed show very similar dynamics throughout learning. **b,** Model with an extra term added in the loss function (cross-entropy loss, CE) to minimize the classification error on direction of motion. The decoding performance of the downstream neuron is higher for the direction of motion. **c,** Model with an additional term on the loss function to limit the activity of the hidden layer in the autoencoder (L2-norm). This configuration of network parameters is similar to Fig. 4b with the downstream neuron reading out from the difference between the reconstructed output and the input. The model discards information about speed and only keeps information about direction of motion. **d,** Same network configuration as [c] with a sparsity penalty that is too large. The network discards information about both speed and direction of motion. Error bars in all panels correspond to SEM across independent simulations (*n* = 50).

**Extended Data Figure 9.**
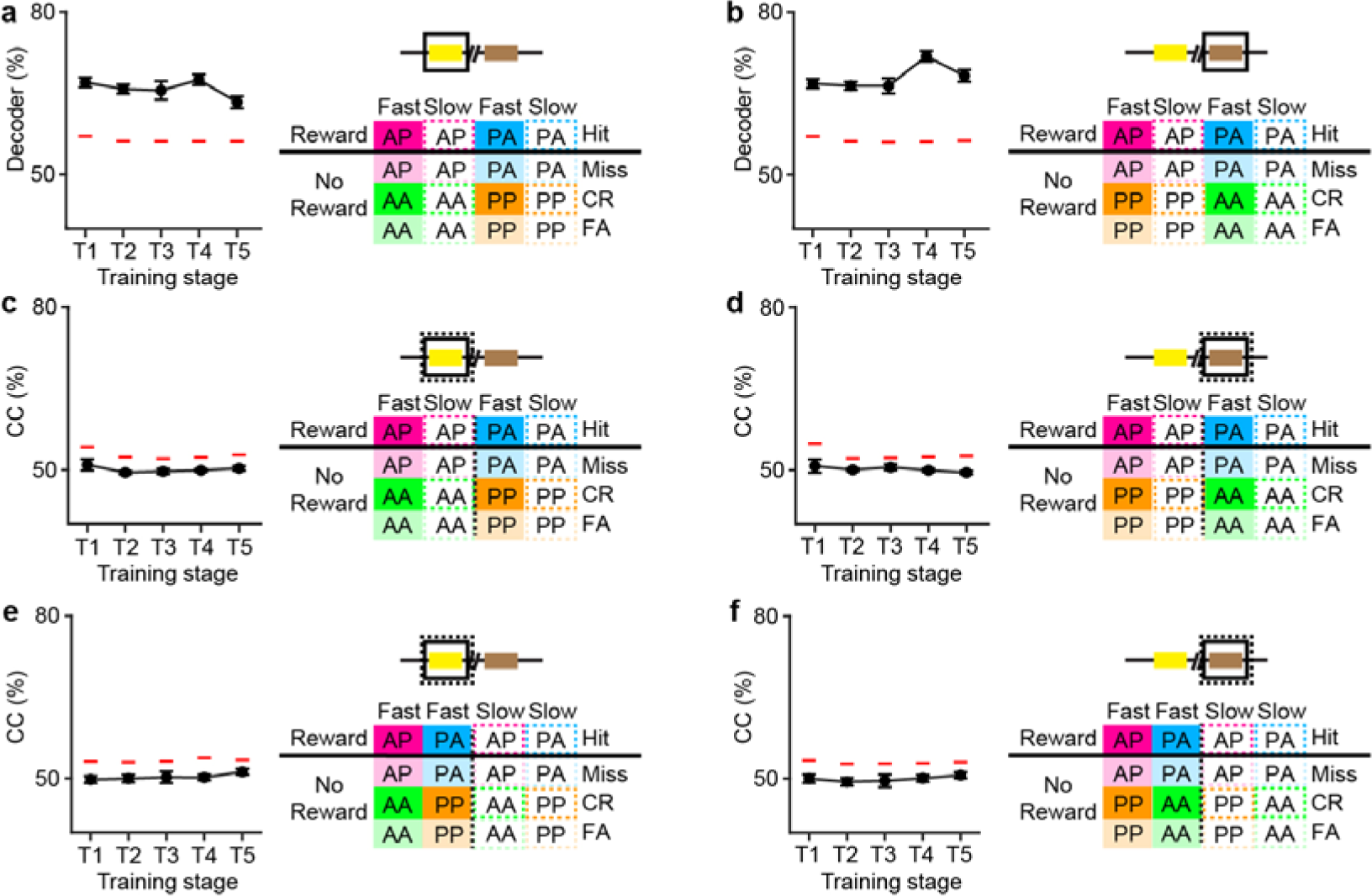
Population coding of reward prediction during sample and test periods and its relationship to stimulus coding. **a**, Linear decoder performance of sample period activity to rewarded conditions across training. **b**, Linear decoder performance of test period activity to rewarded conditions across training. **c**, Cross-condition performance of sample period activity trained to rewarded conditions and tested on stimulus direction conditions across training. **d**, Cross-condition performance of test period activity trained to rewarded conditions and tested on stimulus direction conditions across training. **e**, Cross-condition performance of sample period activity trained to rewarded conditions and tested on stimulus speed conditions across training. f, Cross-condition performance of test period activity trained to rewarded conditions and tested on stimulus speed conditions across training. Error bars = SEM. Red line = 95^th^ percentile performance of the shuffled distribution.

**Extended Data Figure 10.**
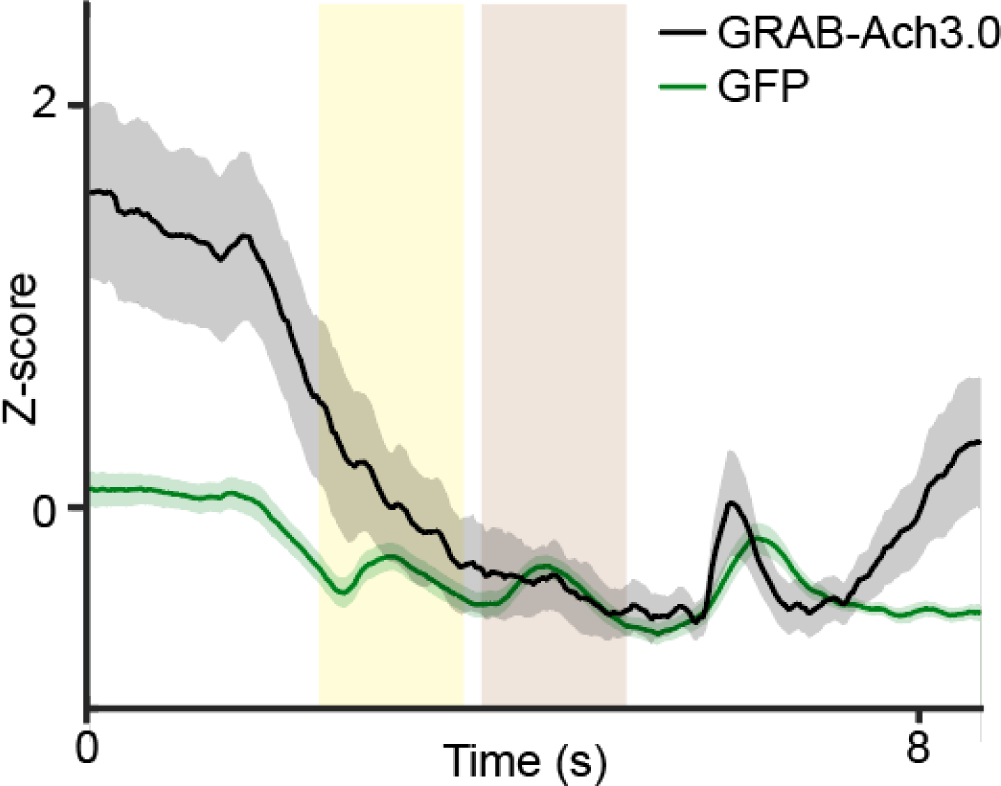
Validation of GRAB-Ach3.0. Z-scored fluorescence traces across the trial period during T2 sessions in task trained animals expressing either GRAB-Ach3.0 or GFP. *n* = 16 T2 sessions from 4 GRAB-Ach3.0 animals, 17 T2 sessions from 2 GFP animals.

**Extended Data Figure 11.**
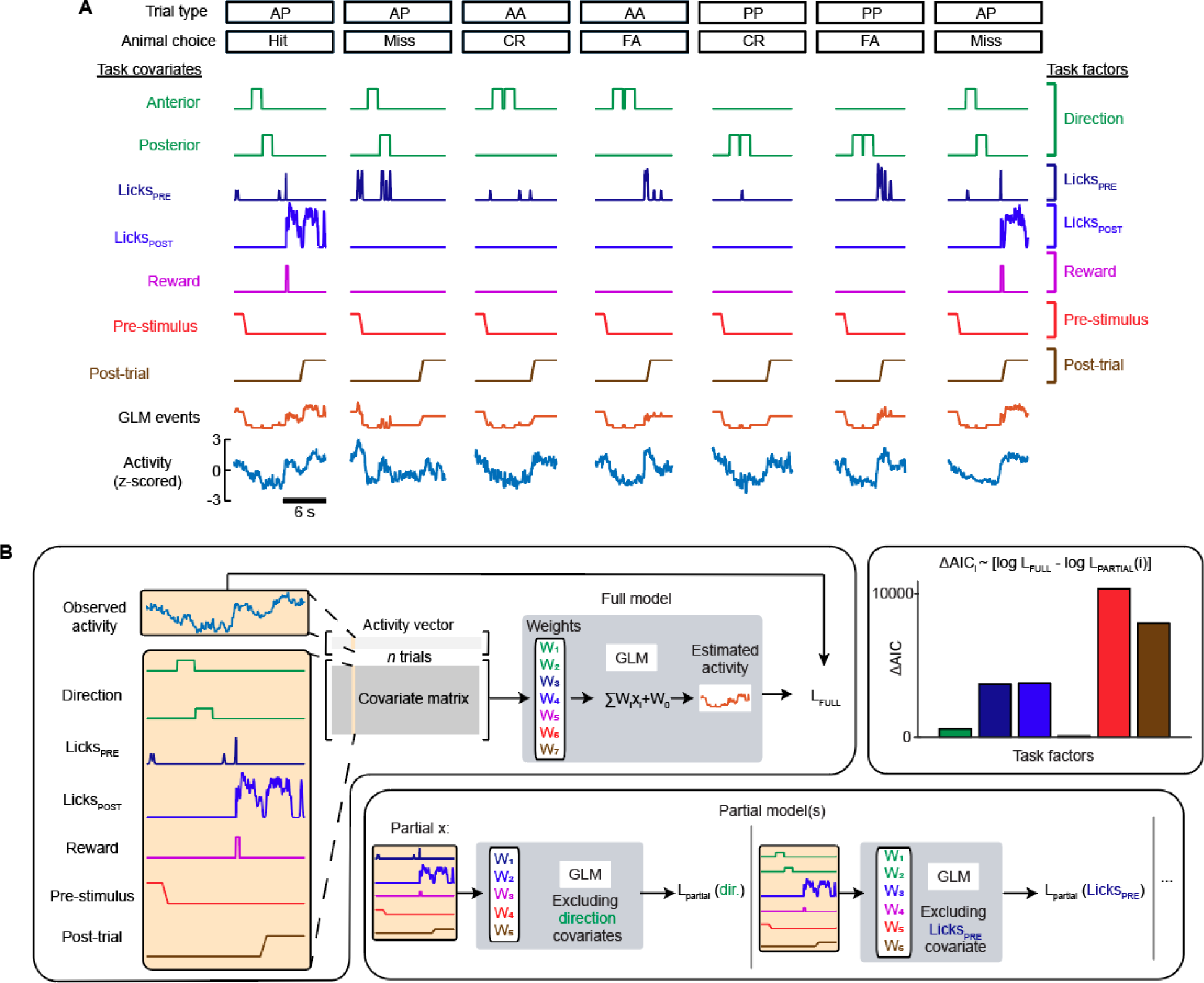
Task encoding of acetylcholine signals. **a,** Overview of covariate representations and their corresponding task factors used in the GLM for acetylcholine signals over six trials. **b,** Schematic of full and partial models used to calculate ΔAIC for individual task factors.

**Extended Data Figure 12.**
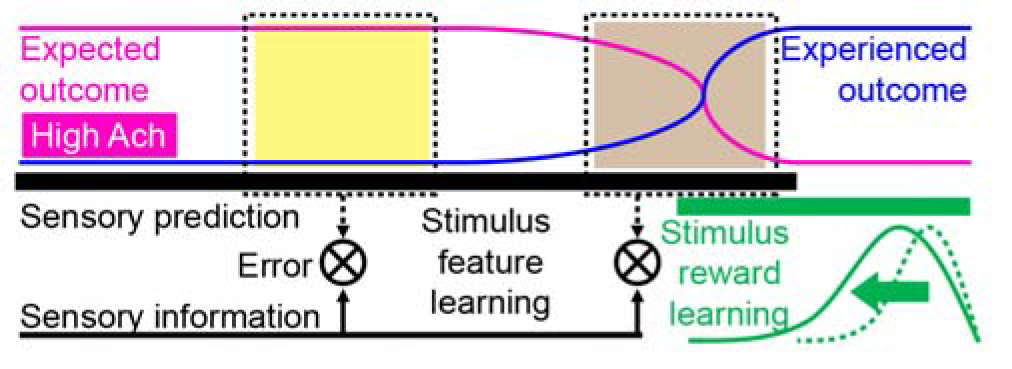
Model of predictive map in Prh. Prh forms a model of task-relevant stimulus information through error learning. Differences in predicted stimulus features elicit sensory prediction errors. Stimuls-reward associations emerge in a retrograde manner from reward outcomes and generalize to similar stimulus-reward contingencies. Expected outcomes are linked to experienced outcomes via a network space that is regulated by cholinergic signaling.

## SUPPLEMENTARY TEXT

### S1. Home-cage task training

In this study, two variations of the tactile working memory task were used to study the role of Prh in abstract sensory learning. To assay the effects of inactivating Prh on behavior, the home cage version of the task was designed to train animals in an unbiased manner. Task training occurred in a training module consisting of a narrow passageway that restricted movement of freely moving animals so that head position was consistent throughout training for reliable delivery of whisker stimulus and water reward (Extended Data Fig. 1a). For whisker stimulus, commercial grade sandpaper (3M; P100) was mounted along the outside edge of a 6 cm diameter rotor, attached to a stepper motor (Zaber) to deflect the whiskers. This was mounted onto a linear stage (Zaber) to place the rotor within whisker reach.

For lick sensing and water delivery, an angled dispensing needle (75165A22; McMaster-Carr) served as a water port. This was attached to a capacitive touch sensor (AT42QT1010; SparkFun) that dispensed 5-7 µL of water through a miniature solenoid valve (LHDA0531115H; The Lee Company). Unlike head-fixed behavior (**Supplementary Text S2**), persistent and impulsive licking was prevalent during freely moving behavior. Attempts to train home-cage animals to learn a go/no go stimulus-reward contingency were not successful due to impulsive licking (data not shown). For these reasons, a two-alternative forced choice (2AFC) task structure using two lick ports was employed for home-cage behavior. To further discourage impulsive licking, lick spouts were mounted onto a linear actuator (L12-P; Actuonix) and only presented to the animals during the report period. This differed from head-fixed training in which lick spouts were fixed always in reach of the animal. Air puffs were controlled using a 12V solenoid (EV-2-12; Clippard). Task training was performed using a custom written LabVIEW software (National Instruments) to control hardware and a data acquisition interface (USB-6008; National Instruments) for measuring licks, water delivery, and air puff delivery.

The task was designed for live-in conditions in which trials were self-initiated and task parameters automatically adjusted based on performance. A single training module was connected to three cages, each containing a singly-housed mouse. Mice were singly-housed to avoid social interactions that would interfere with equal access to task training. Head-fixed mice were similarly singly-housed to minimize potential damage to their implants. Cages were connected via passageways to a common meeting chamber. For each passageway, access to the training module via the meeting chamber was regulated by mechanical doors. These doors were controlled by servos operated by an Arduino microcontroller. Door closing was trigged by an infrared beam break sensor placed between the door and home cage in order to ensure that the door did not close while the animal was in the training module.

Access to home-cage training was scheduled similarly to head-fixed task conditions to ensure equivalent water deprivation periods, motivation levels, session duration, and trial numbers.

Each animal gained daily access to the training module for two, two-hour sessions (Extended Data Fig. 1b). To ensure that each animal performed the task across all dark portions of light-dark cycle, the scheduled animal order was rotated daily. At the end of each session, the training module break beam sensor was deactivated to prevent trial initiation. A continuous train of air puffs was delivered into the chamber signaling the animal to exit and for the door to close behind them (Extended Data Fig. 1c). A USB radio frequency identification (RFID) reader above the meeting chamber was used to ensure that the correct animal accessed the training module at the properly scheduled time.

### S2. Head-fixed task training

The head-fixed version of the task was designed to reliably image neuronal activity during learning under highly consistent, well-controlled stimulus conditions. A go/no go stimulus-reward contingency was employed to characterize activity patterns related to stimulus information with and without reward associations. Similar to home-cage training, the task was performed using a custom written LabVIEW software (National Instruments) to control hardware and a data acquisition interface (USB-6008; National Instruments) for measuring licks, water delivery, and air puff delivery. A water port was attached to a capacitive lick sensor (AT42QT1010; SparkFun) that dispenses 5 to 6 µL of water through a miniature solenoid valve (0127; Buekert). For the rotation stimulus, commercial grade sandpaper (3M; roughness: P100) was mounted along the outside edge of a 6 cm diameter rotor, attached to a stepper motor (Zaber) to deflect the whiskers which was mounted onto a linear stage (Zaber) to place the rotor within whisker reach.

Given the time demands of the experiment for operating the two-photon microscope through learning (∼70 sessions, 2 sessions per day, 7 days per week), ensuring successful training was a priority for animals undergoing imaging. Given the natural variability in learning across individual animals, experimenters manually adjusted a range of behavioral parameters designed to reinforce correct choice behavior (**Supplementary Text S4**).

### S3. Training stages

The task settings defining each training stage in the home cage (Table 1) and head-fixed (Table 2) training task were largely similar with the following exceptions. For the head-fixed task, the proportion of non-match versus match trials were gradually changed from 0.9/0.1 to 0.5/0.5 (non-match/match) over the course of the first 5 T1 sessions. The purpose of this was to acclimate the animals to licking for reward and to avoid miss trials by providing a high proportion of rewarded (non-match) stimulus trials and gradually exposing animals to the non-rewarded (match) stimulus trials. For the home cage task, the proportion of non-match versus match trials were set to 0.5/0.5. During early T1 sessions, the maximum consecutive trials belonging to either match or non-match stimulus was set to 1. This meant that water reward alternated between each lick port in order to acclimate the animal to licking to each port. The target spout alternated between trials through four sub-stages which taught animals how to receive rewards and gradually introduced the moving parts of the task. In the first sub-stage, the texture was positioned against the training module but did not provide directional stimuli. Instead, animals were able to trigger a trial and lick when an audible tone was played in order to receive a water reward. With consistent lick responses, the delay between triggering a trial and the tone indicating the report period was increased from 100ms to 6s, approximating the time course of a trial with two stimuli and a 2s delay. In the second sub-stage, the sample and test stimuli were presented and the report period was still indicated with a tone. This tone was removed during the third sub-stage. The fourth sub-stage introduced linear movement of the texture, withdrawing it at the end of a trial and moving it to presentation position for the sample and test periods. The maximum number of consecutive trials with the same target spout was then increased to 3 in the fifth sub-stage to randomize the stimulus conditions.

During T4, the delay between the sample and test stimulus was gradually increased through a progression of sub-stages. An initial delay was used at the beginning of the session. Behavioral performance was measured every 15 trials. The delay was increased by a defined increment if performance exceeded >85% correct (*d’* > 2.07) over the past 15 trials up to a maximum of 2 seconds. If the overall performance for the session was *d’*>1.68, the animal advanced to the next T4 sub-stage in which the starting delay and increment was greater than that used in previous session. The rotor was withdrawn once animals could begin sessions with delays of 2 seconds. In general, head-fixed animals could readily adapt to the rotor withdrawal during the delay period. Initial piloting of the same training progression during home-cage task training suggested that animals had difficulty with adapting to this transition. For this reason, the training protocol in the home-cage task was modified to include a gradual withdrawal of the rotor occurring concurrently with the gradual increase in delay period.

During T5, delays were randomly varied between 2, 3, and 4 seconds for head-fixed animals to examine sequential activity across varying delay periods. In home-cage animals, the delay was fixed at 2 seconds. Finally, slow speed stimulus conditions were included for head-fixed task in order to measure activity related to relevant and irrelevant stimulus features but were not included during the home-cage task since the motivation of the latter was to broadly assay the dependence of Prh on task learning.

### S4. Reinforcing correct choice

Due to the complexity of task conditions and stimuli, we observed that individual animals adopted a range of incorrect choice strategies early during task training. Occasionally, behavioral lapses were also observed in which animals demonstrated correct choice strategies across extended trial periods but then reverted to incorrect choice strategies. Incorrect choice strategies were categorized as report bias, primacy bias, and recency bias. A set of task parameters were included in the training protocol to identify and correct for these biases without changing the stimulus-reward contingency (Table 3).

For go/no-go behavior under head-fixed conditions, a report bias was defined as persistent licking of the lick port regardless of stimulus condition. For 2AFC version of the task used in the home cage training system, persistent licking of one of two lick ports regardless of stimulus condition was considered a report bias. Report bias primarily contributed to poor task performance early in training during T1 and was also occasionally observed at the beginning of behavior sessions in trained mice. For the go/no go head-fixed task, report biases were defined by a high fraction of total hit and false alarm trials. For 2AFC home cage task, report biases were defined a high fraction of hit and false alarm trials attributed to one of the two lick ports. Depending on the severity of the report bias, two corrective strategies were adopted. The first strategy is the use of punishment to discourage licking of the incorrect stimulus condition. Punishment consisted of a combination of time out and air puffs to the face. Initially introduced punishment was mild and gradually became more severe with longer time outs and multiple air puffs considered as more severe punishment. Tolerance for punishment can vary for individual animals (data not shown). For both task conditions, animals disengaged from the task if punishment was too aversive, resulting in miss trials. Punishment levels are reduced if misses increase. In addition to adjusting punishment levels, the probability of stimulus conditions was also adjusted to increase the frequency of the incorrect stimulus condition in order for animals to “practice” the correct response. Typically, non-match and match stimulus conditions were presented at 50% probability. This was increased up to 80% for the incorrect stimulus condition depending on the severity of the report bias.

A primacy stimulus bias represented incorrect choice strategies in which the animal responded based on whether the sample stimulus was A or P. In contrast, a recency stimulus bias represented incorrect choice strategies in which the animal responded based on whether the test stimulus was A or P. These biases were operationally defined as differences in performance between the two stimulus conditions belonging to the same category (AP vs. PA for non-match, AA vs. PP for match). Typically for each stimulus category, one of the two possible stimulus conditions is presented with 50% probability with respect to the other. To correct for primary or recency bias, the probability of stimulus conditions belonging to the same category was adjusted to increase the frequency of the incorrect stimulus condition in order for animals to “practice” the correct response.

